# Large-scale multi-omics analyses identified root-microbiome associations underlying plant nitrogen nutrition

**DOI:** 10.1101/2024.02.05.578621

**Authors:** Nannan Li, Guoliang Li, Danning Wang, Lige Ma, Xiaofang Huang, Zhen Bai, Yongfeng Wang, Meng Luo, Yu Luo, Yantao Zhu, Xulv Cao, Qirui Feng, Ying Xu, Jianxin Mu, Ran An, Cuiling Yang, Hao Chen, Xiaodan Li, Yachen Dong, Jianhua Zhao, Lixi Jiang, Yong Jiang, Jochen C. Reif, Frank Hochholdinger, Xinping Chen, Daojie Wang, Yanfeng Zhang, Yang Bai, Peng Yu

**Affiliations:** College of Resources and Environment, and Academy of Agricultural Sciences, Southwest University, Chongqing, 400715, China; Emmy Noether Group Root Functional Biology, Institute of Crop Science and Resource Conservation (INRES), University of Bonn, Bonn, Germany; Crop Functional Genomics, Institute of Crop Science and Resource Conservation (INRES), University of Bonn, Bonn, Germany; Leibniz Institute of Plant Genetics and Crop Plant Research (IPK), Stadt Seeland, Germany; Peking-Tsinghua Center for Life Sciences, College of Life Sciences, Peking University, Beijing 100871, China; Yazhouwan National Laboratory, Sanya, 572025, China; Hybrid Rapeseed Research Center of Shaanxi Province, Yangling, Shaanxi 712100, PR China; Institute of Crop Science, Zhejiang University, Hangzhou 310058, PR China; College of Agriculture, State Key Laboratory of Crop Stress Adaptation and Improvement, Henan University, Kaifeng 475004, Henan, China; Shanghai Majorbio Research Institute, Shanghai, 201203, PR China; Interdisciplinary Research Center for Agriculture Green Development in Yangtze River Basin, Southwest University, Chongqing 400715, China; Research Center for Intelligent Computing Platforms, Zhejiang Lab, Hangzhou, 310012, PR China

**Author notes:** To whom correspondence should be addressed: <u>Nannan Li Daojie Wang Yanfeng Zhang Yang Bai, Peng Yu</u> (lead contact). These authors equally contributed to this work (Nannan Li, Guoliang Li, Danning Wang, Lige Ma, Xiaofang Huang, Zhen Bai). **Author contributions** N.L. and P.Y. designed the study; P.Y. coordinated the whole project; P.Y. and G.L. performed the population genetic and genomic prediction analysis; D.W. analysed the microbiome data; G.L. and D.W. performed all statistical analysis; L.M., X.H. and Z.B. jointly performed bacterial cultivation, inoculation and functional validation experiments; N.L., Y.L., Z.B., Y.Z., X.C., X.J., Y.X., J.M., R.A., C.Y. and H.C. performed the field sampling; N.L., Y.L., Z.B. and Y.Z. conducted the plant nutrient analyses. J.C.R., F.H., X. C., D.W. and Y.Z. interpreted the data. P.Y. drafted the manuscript and all authors read and approved the final version of the manuscript.

**Keywords:** *Brassica napus*, eQTL, GWAS, ionome, rhizosphere microbiome, root transcriptome

## Abstract

The microbiome determines the performance and fitness of the host plant. Nevertheless, the causal interaction between host genetic variation, gene regulation and the impact of the microbiome on the host phenotype remain unknown. Here we generated 1,341 paired root transcriptome, rhizosphere microbiome and root ionome samples and performed a multi-omics analyses of the host-microbe association at the root-soil interface using 175 rapeseeds (*Brassica napus* L.) resequenced ecotypes at two field environments. We observed the highest statistically explained variance for root nitrogen uptake among natural ionomic variation by overall transcriptome-wide gene expression and microbial abundance variation. Moreover, we identified significant genome-wide associations for 203 highly heritable amplicon sequence variants (ASVs) at multiple genetic loci regulated by eQTL hotspots associated with nitrogen metabolism components. These associations involved a central bacterial genus (*Sphingopyxis*), which plays a dominant role on gene regulatory effect on its variation regulated by eQTL hotspots. In addition, we performed high-throughput bacterial cultivation from rapeseed roots and subjected *Sphingopyxis* to whole genome sequencing. Finally, targeted metabolite profiling and confocal imaging assays demonstrated a host-microbiome regulatory effect on *Sphingopyxis* established by lateral root development and plant nitrogen nutrition. In summary, our integrative approach reveals the genetic basis of host-microbiome trait associations in the transcriptional, nutritional and environmental domains and suggests that the microbiome might have causal effects on root development with implications towards the breeding of nutrient-efficient crops.

## Introduction

The rhizosphere, which is the soil region in the vicinity of plant roots, represents a unique environment in which complex biological interactions between the plant root system and the entity of soil microbes designated as the rhizosphere microbiome (Philippot et al., 2013) take place. This microbiome influences root traits and functions affecting plant health (Durán et al., 2018), assists with host nutrient homeostasis (Salas-González et al., 2020), protects against biotic and abiotic stresses (Cheng et al., 2019), alters the plant developmental program (Finkel et al., 2020) and acts as important driver of ecosystem functioning (Banerjee et al., 2018). Microbes inhabiting plant roots and its affected rhizosphere harbor a wealth of functional traits such as metabolic properties (Hu et al., 2018; Zhalnina et al. 2018; Huang et al. 2019; Wilschut et al., 2019) and immune system functions (Lebeis et al., 2015; Carrión et al., 2019). Indeed, small host-mediated changes in a microbiome can have large effects on host fitness (Haney et al., 2015), although the effect of host genotype is minor in comparison with soil edaphic factors (Bulgarelli et al., 2012; Lundberg et al., 2012). Genome-wide association studies (GWAS) are an efficient approach for identifying the genetic variation associated with microbes in a community context (Beilsmith et al., 2019). Quantitative analyses of microbiome community composition and genetic plant diversity identified heritable microbes and allowed to integrate them as suitable phenotypes into GWAS (Horton et al., 2014; Cordovez et al., 2019; Deng et al., 2021; Escudero-Martinez et al., 2022; Meier et al., 2022; Oyserman et al., 2022; Wang et al., 2022). Nevertheless, a clear understanding of how the rhizosphere microbiome forms and how its function is regulated by host gene expression beyond the influence of plant fitness and agroecosystem functions in crop species remains elusive.

Nitrogen is the quantitatively most important macro mineral element for plants and plays a critical role for crop yield. Excessive use of nitrogen fertilizers in agriculture are a global threat to ecosystems and human health (Guo et al., 2005; Liu et al., 2010). In the past decade, nitrogen fertilizer application into terrestrial soils contributed to ∼60% of the total increase of atmospheric nitrous oxide emission (Tian et al., 2018; Chen et al., 2014). Essentially all mineral nutrients consumed by humans are captured by elaborate root systems and transported through the rhizosphere (Marschner, 2012). Numerous examples have demonstrated that beneficial symbiotic associations of roots and the rhizosphere with microbes can improve plant performance by improved nutrient uptake, tolerance to abiotic stresses and avoidance of soil pathogens under nutrient stress conditions (Mendes et al., 2013; Yu et al., 2016). Increasing genetic evidences demonstrate that the root microbiome drives the assemblage of microbial communities based on nutritional properties such as acquisition of nitrogen (Zhang et al., 2019a; Yu et al., 2021; He et al., 2023) and direct sensing of the phosphate stress (Castrillo et al., 2017) from the soil.

In this context, it is of fundamental importance to understand the genetic basis and the regulatory variation of host–microbiome associations and the mechanisms underlying the assemblage and function of the crop microbiome that affect plant nutrition. The present study incorporates integrative rhizosphere microbiome, host genome-wide and transcriptome-wide association analyses in two independent environments that quantify interactions among genotypes, environment and plant nutritional status to predict the fitness and health in rapeseed. To this end, beneficial associations between the soil microbiome and the root system will pave the way for designing crop varieties with ideal root phenotypes to recruit and activate microbiomes with broad benefits for enhancing crop productivity, efficient nutrient acquisition and agroecosystem resilience.

## Results

### Integrative multi-omics rapeseed root transcriptome and rhizosphere microbiome

Overall, we aimed to understand the reciprocal associations between genetic variation, root gene regulation and rhizosphere microbiome assembly, and how such associations influence the plant nutritional phenotype in rapeseed (Fig. 1a). We subjected 175 genetically divers and fully sequenced rapeseed lines (Supplemental Fig. 1a; Supplemental Dataset 1) covering spring, winter and semi-winter ecotypes to plant-microbe interactions under two field conditions in China (Supplemental Fig. 1b). To consistently compare all genotypes, the lateral roots along the longitudinal region of the root tap at the flowering stage were dissected and pooled. Half of these pooled samples were immediately washed and dried with clean tissue as the root sample for RNA sequencing analysis and the other half with soil closely attached to these lateral roots was defined as the rhizosphere sample. In addition, we collected bulk soil from the unplanted plots. The isolation and extraction of rhizosphere samples was performed according to our previous work (Yu et al., 2021). We used RNA sequencing and 16S (V4) rRNA gene sequencing to characterize root gene expression and the rhizosphere microbiome composition, respectively. A principle component analysis (PCA) demonstrated that location explains most of the variance for both, root gene expression and rhizosphere microbiome, while the genotype explained more variance on gene expression than the microbiome (Supplemental Figs. 2 and 3).

**Figure 1.**
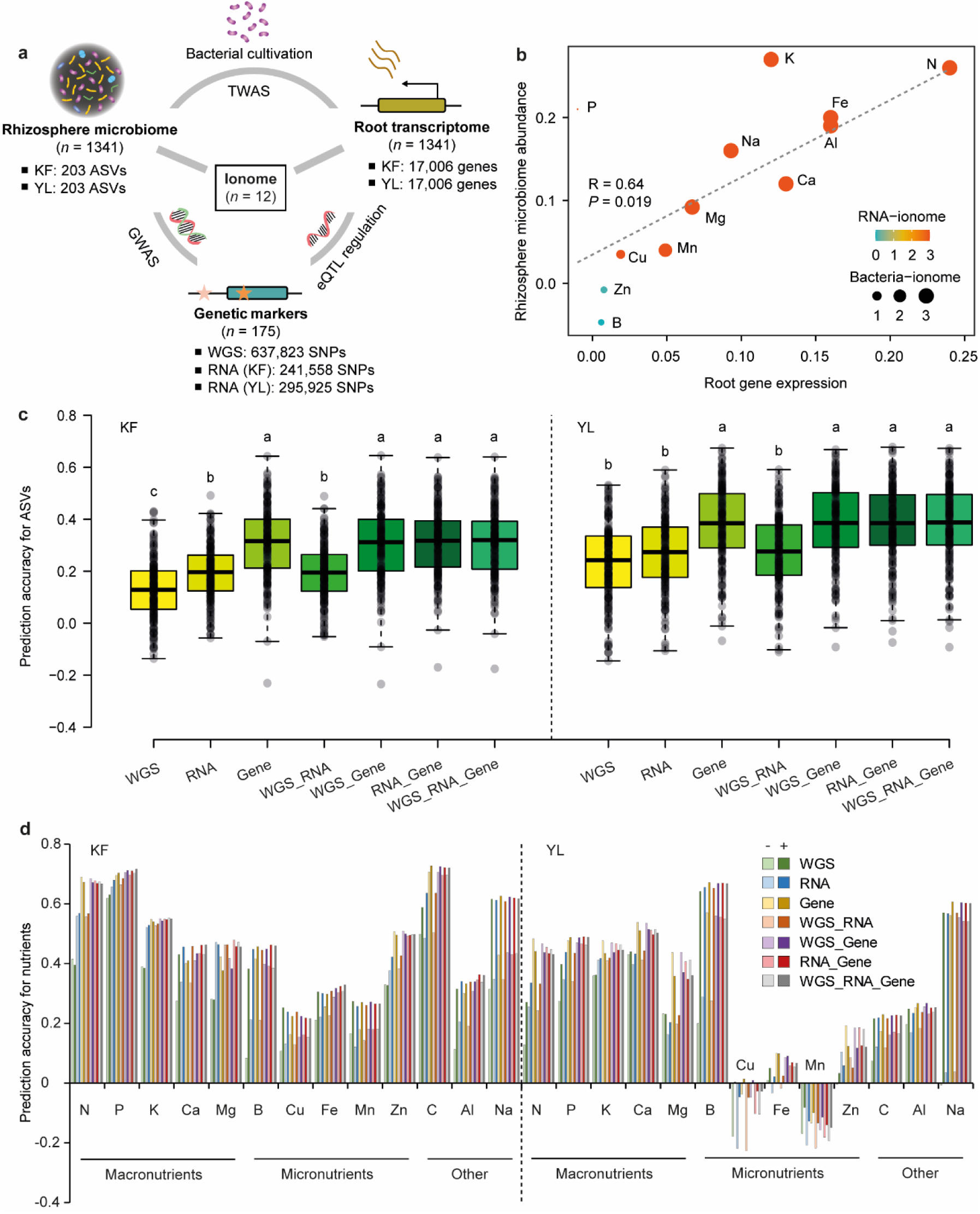
Genomic and transcriptomic analyses of host-microbiome association and plant nutritional traits in *B. napus*. **a**, A schematic illustration of omics datasets used in this study. KF, Kaifeng; YL, Yangling. ASV, amplicon sequence variant; SNP, single-nucleotide polymorphism. GWAS, genome-wide association studies; TWAS, transcriptome-wide association studies; eQTL, expression quantitative trait loci; Whole genome sequencing (WGS) data was derived from Wu et al., 2019. SNP data were extracted from the RNA sequencing data for both KF and YL locations. Ionome data includes the concentration of 12 different mineral nutrients. **b**, Spearman correlation between gene expression, rhizosphere microbiome diversity and plant ionomic traits. Different color codes and sizes of the dots indicate the significance (−log_10_) of pair-wise correlations between gene expression or microbiome diversity and ionomic traits. **c,** Different scenarios for predicting abundance of bacterial ASVs of the rhizosphere microbiome. The box plot shows the prediction ability of 203 ASVs based on different omics data. **d**, Plant nutritional trait prediction using genetic, transcriptomic information combined with microbiome feature. The bar plot shows the prediction ability of 13 mineral elements based on WGS, RNA, gene data with or without ASVs information. The results show that the ASVs information does help to improve the prediction ability of 13 mineral elements.

We next applied WGCNA to cluster all expressed genes and identified 35 gene co-expressed modules across all genotypes at two locations (Supplemental Dataset 2). For the microbiome feature, we filtered and normalized all ASVs to identify the highly abundant and confident ones at the family and genus levels (Supplemental Dataset 3). Moreover, we calculated the broad sense heritability (*H^2^*) and selected in total 203 highly heritable ASVs a *H^2^* >0.15 (Walters et al., 2018) accordingly (Supplemental Dataset 4). We further integrated the gene co-expressed modules with the highly heritable ASVs, relative abundance of different families and genera as phenotypic traits (Supplemental Dataset 5). Interestingly, we identified four co-regulated modules (darkorange, lightcayan, lightsteelblue1 and royalblue) which are highly associated with different microbial traits (ASV, families and genera) (Supplemental Fig. 4a). We functionally characterized these modules in gene ontology (GO) categories and demonstrated that in total 663 unique GO terms were enriched in the biological pathways (Supplemental Fig. 4b). In particular, several representative biological pathways such as “carbon and nitrogen biosynthesis and metabolism”, “regulation of growth and metabolism”, “abiotic and biotic responses”, “embryonic and postembryonic root development”, “transport” and “organization” were identified by REVIGO (Supplemental Fig. 4c; See Methods). Moreover, we identified several dominant bacterial families such as Bacillaceae, Chitinophagaceae, Comamonadaceae, Gemmatimonadaceae, Nocardioidaceae, Pseudonocardiaceae, Rhizobiaceae, Sphingomonadaceae, Vicinamibacteraceae, Xanthobacteraceae and Xanthomonadaceae across all these heritable ASVs (Supplemental Dataset 4). Taken together, these functional pathways and dominant bacterial taxa might be important indicators for linking plant phenotypic traits.

### Overall genomic, transcriptomic and microbial predictions for plant nutritional phenotypes

We investigated the nutritional status and variation in the ionomic compositions (N, P, K, Ca, Mg, Fe, Mn, B, Cu, Zn, Al and Na) of the root at flowering stage for the 175 rapeseeds accessions grown at two locations. We first performed Mantel’s statistical test and identified a significantly positive correlation (R = 0.64, *P* = 0.019) between root gene expression and rhizosphere microbiome abundance across different ionomic traits (Fig. 1b). In detail, the highest correlation was identified between gene expression and concentration of N (R = 0.23, *P* = 0.001), followed by Fe (R = 0.16, *P* = 0.001), Al (R = 0.16, *P* = 0.001), Ca (R = 0.13, *P* = 0.001) and K (R = 0.12, *P* = 0.001) (Fig. 1b). A similar correlation pattern was found between the rhizosphere microbiome and ionomic concentrations, i.e. K (R = 0.27, *P* = 0.001), N (R = 0.26, *P* = 0.001), P (R = 0.21, *P* = 0.001), Fe (R = 0.2, *P* = 0.001) and Al (R = 0.19, *P* = 0.001) (Fig. 1b).

To understand how genetics and gene regulation affect microbial abundance, we next applied different scenarios of genomic predictions on the abundance of microbial ASVs using genomic markers, transcribed markers, gene expression data and different combined models (Supplemental Fig. 5; See Methods). Interestingly, gene expression data alone showed the best prediction accuracy (KF: 31%, *P* <3.37e^−06^; YL: 38%, *P* <5.17e^−11^) for microbial ASVs compared to other scenarios. A combination of gene expression with genetic markers did not generate an additional advantage for predicting microbial ASVs at both field locations (Fig. 1c). We further assessed the genomic prediction ability of different ionomic traits using the rhizosphere microbiome alone or in combination with different prediction scenarios at both fields. The combination of plant genomic, transcriptomic and rhizosphere bacterial community composition provided the highest average prediction ability (KF: 53%; YL: 40%) for macronutrients (N, P, K, Ca and Mg) and the lowest prediction (KF: 30%; YL: 10%) for micronutrients (B, Cu, Fe, Mn and Zn) (Fig. 1d). Based on these results, we hypothesize that the gene regulatory effect might be more influential in determining microbiome composition than genetic markers. Under our field conditions, genomic predictions were better for macronutrient ionomic traits than for micronutrients. Thus, integration of root transcriptional information with the rhizosphere bacterial microbiota will largely advance our understanding of the structure and function of the rhizosphere microbiome underlying macronutrients acquisition in rapeseed.

### Genome-wide and transcriptome-wide association analyses for microbial ASVs

To systemically understand the genetic basis and gene regulatory effect on microbial assembly, we integrated GWAS and TWAS for 203 highly heritable ASVs with plant ionomic traits. We selected 345,289 variations in 175 *B. napus* accessions for GWAS for ASVs at 2 locations, respectively. An FDR correction of *p* <0.05 has been employed as the genome-wide threshold for 203 ASVs. Moreover, we detected 169 marker-microbial trait associations for 34 ASVs, which traced back to 99 independent QTLs, in KF. Furthermore, we observed 510 marker-microbial trait associations for 59 ASVs in YL, which traced back to 317 independent QTLs (Figure 2a). Within the QTL interval, we found 184 candidate genes in KF and 309 in YL (Supplemental Dataset 6). There were 14 common ASVs (“ASV340”, “ASV260”, “ASV339”, “ASV704”, “ASV235”, “ASV1045”, “ASV435”, “ASV1580”, “ASV1163”, “ASV198”, “ASV106”, “ASV46”, “ASV478”, “ASV1178”) that detected QTLs in both KF and YL. Four QTLs were detected in both KF and YL. For instance, the QTL represented by lead SNP A07_5345677 was detected for ASV1646 in KF and for ASV3776, ASV478, ASV85 and ASV767 in YL. The QTL from 8,243,938 to 8,244,269 in chromosome A02 was detected for ASV1580 in KF and for ASV1603 in YL, the QTL from 28,696,661 to 28,696,967 in C07 was detected for ASV106 in KF and for ASV1603 in YL, the QTL from 31,122,762 to 31,122,769 in C07 was detected for ASV478 in KF and for ASV1592 in YL.

**Figure 2.**
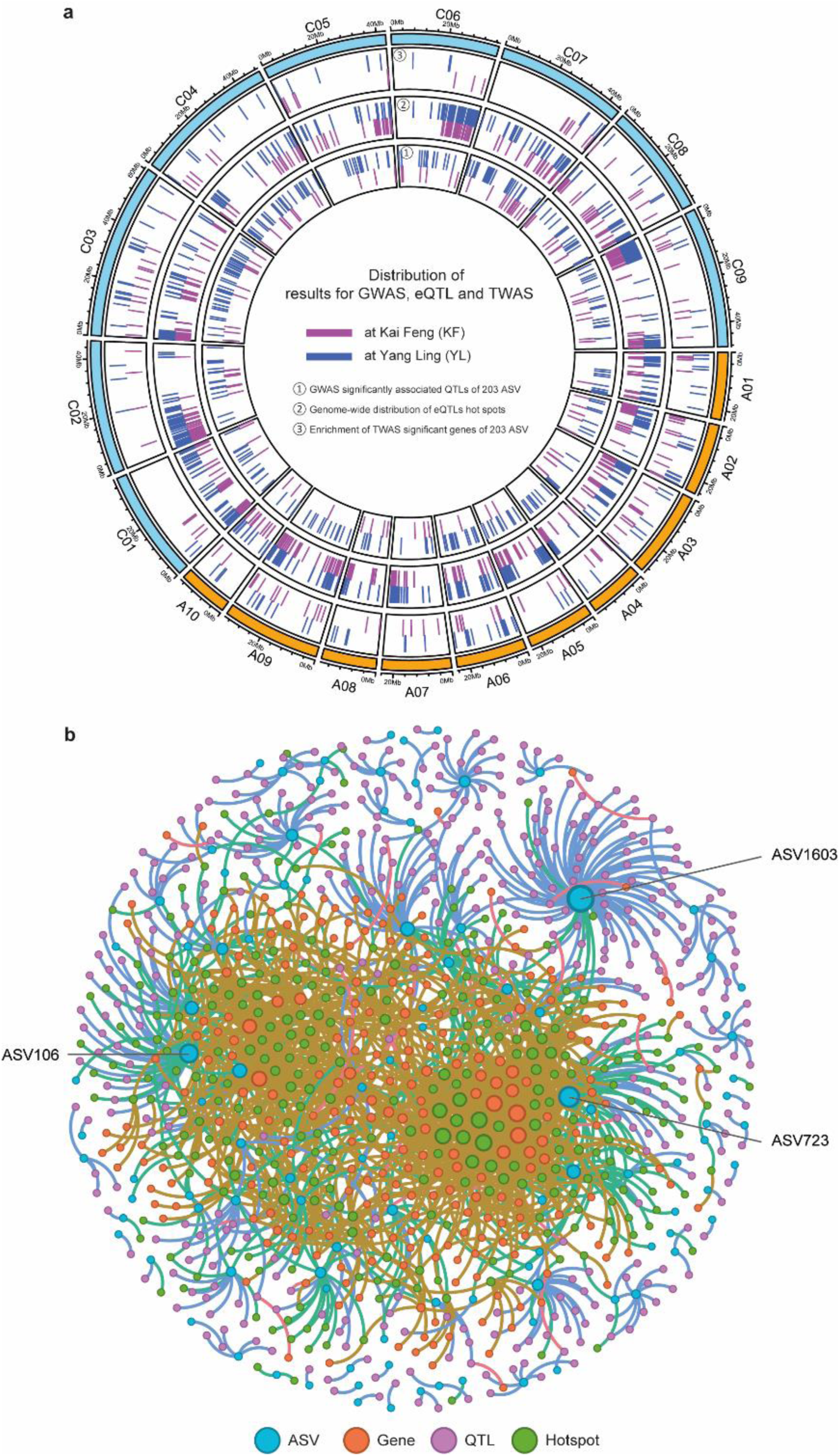
Integrative genomic and transcriptomic analyses of host-microbe associations in B. napus. **a**, The genomic distribution of QTLs, eQTL and the enrichment of TWAS significant genes of all 203 ASVs. From inner to outer: ①GWAS significantly associated QTLs of 203 ASVs, ②eQTL hotspot detected by eGWAS, and ③TWAS significant genes identification. And the pink and dark blue indicate the results in KF and YL. **b**, The ASV-GWAS-eGWAS-TWAS regulated network integrates results from KF and YL. Blue circles den ote 106 amplicon sequence variants (ASVs) with significant genes or QTLs, and the three ASVs with mos t significant gene and QTL are ASV1603, ASV723 and ASV106. Orange circles represent 287 TWAS significant genes of ASVs. Purple circles indicate 389 GWAS significantly associated QTLs of ASV. Green cir cles represent 250 eQTL hotspots, and the shape size indicating the number of regulatory genes.

Transcriptome-wide association studies (TWAS) integrate genome-wide association studies (GWAS) and gene expression datasets to identify gene–trait associations. We selected 17,006 gene expressions in 175 *B. napus* accessions for TWAS for ASVs at 2 locations, Kaifeng (KF) and Yangling (YL), respectively. An FDR correction of *p* <0.05 has been employed as the genome-wide threshold for 203 ASVs. Overall, we detected 174 gene-microbial trait associations (154 unique genes) for 36 ASVs in KF; and 181 gene-microbial trait associations (136 unique genes) for 46 ASVs in YL (Supplemental Dataset 7).

### eQTL regulation of heritable ASVs associated with nitrogen genes

Gene expression functions as a bridge between the genotype and the phenotype. To understand the contribution of gene expression in the variation of microbial ASVs abundance among rapeseed plants with different genotypes, we performed root RNA sequencing of a core collection comprising 175 *B. napus* accessions in Kaifeng (KF) and Yangling (YL), respectively. In total, eGWAS identified 79,810 eQTLs associated with the expression of 12,293 genes (eGene regulated by eQTL) in KF, and 121,955 eQTLs associated with the expression of 12,322 genes in YL (Supplemental Dataset 8). These eSNP located in eQTLs and eGenes were from all 19 chromosomes (Supplemental Fig. 6). We found that the associations of eQTLs and eGene located on the same chromosome had a higher significance than those located on different chromosomes (two-sided Wilcoxon rank sum test, *p*-value <2.2×10^-16^; Supplemental Fig. 7), both in KF and YL. For the inter-chromosomal associations, the co-localization of distant eQTL and target genes in the syntenic regions of homoeologous chromosomes (between A-subgenome and C-subgenome) were prevalent (Supplemental Fig. 8). Based on these eQTL results, we detected 317 hotspots in KF and 380 in YL, in which the associations between the hotspot and other chromosomes were enriched, as the vertical lines indicated in Supplemental Fig. 9.

Based on the distance between eQTLs and eGenes, we categorized all eQTLs into 3,421 local eQTLs (<50kb) and 76,389 distant eQTLs (>50kb or in different chromosomes) in KF, and 4,720 local eQTLs and 117,235 distant eQTLs in YL. We found that local eQTLs had a larger effect on expression variation than did distant eQTLs (two-sided Wilcoxon rank sum test, *p*-value <2.2×10^-16^; Supplemental Fig. 8), both in KF and YL. Of the total eQTLs in KF, local eQTLs account for 4.3%, while distal eQTLs account for 95.7%, of which 14.4% occurred on the same chromosome (distant_intraChr) and the other 81.3% were found on different chromosomes (distant_interChr). Of the total eQTLs in YL, local eQTLs account for 3.9%, while distal eQTL account for 96.1%, of which 18.1% occurred on the same chromosome (distant_intraChr) and the other 78% were found on different chromosomes (distant_interChr). Of the eGenes in KF, we found that 3,304 were regulated by local eQTLs and 11,978 by distal eQTLs, and one-third of eGenes (3792, 30.8%) were regulated by only one eQTL or two eQTLs (Supplemental Fig. 9). Of the eGenes in YL, we found that 4,566 were regulated by local eQTLs and 11,976 by distal eQTLs. Overall, 22.8% eGenes (2814) were regulated by only one eQTL or two eQTLs (Supplemental Fig. 9).

#### Construction of the ASV-GWAS-eGWAS-TWAS regulated network

To better understand the interactions between microbial ASVs and the host genotype of *B. napus*, we integrated the results of eGWAS, GWAS, and TWAS to construct a comprehensive multi-omics eQTL regulatory network (Supplemental Dataset 9). Totally, in KF and YL, the network contained 106 ASVs, 389 GWAS significantly associated QTLs of ASV, 287 TWAS significant genes of ASV, and 250 eQTL hotspots, indicating that the genetic and gene regulatory effect on host-microbe association is extremely complex in *B. napus* (Supplemental Dataset 9).

The difference and diversity among ASVs could be better understood through the phylogenetic trees of 203 ASVs, which describe both the evolutionary process and community diversity (Supplemental Fig. 10). To better understand the impact of ASVs, for each gene, a stepwise regression method was used to screen the ASVs affecting gene expression. Then, several influential ASVs would be selected for each gene. Finally, as output, an ASV count table, which reported the number of times each ASV has been selected for all 17,006 genes. The top 10 ASV in KF are ASV23, ASV460, ASV9, ASV179, ASV965, ASV193, ASV269, ASV1750, ASV26, ASV407, and ASV205, ASV5542, ASV344, ASV1766, ASV287, ASV95, ASV38, ASV1307, ASV1565, ASV827 in YL. The ASVs located in the branches including both the top 10 ASVs of KF and YL, of the phylogenetic trees will be the focus of our subsequent research. The only branch that meets this condition is the one which represented by ASV407, ASV723, ASV3382 and ASV5542.

#### The bacterial taxon *Sphingopyxis* supports root development and nitrogen uptake under stress

Among all detected genetic loci with highly abundant bacterial taxa, we selected the bacterial taxon *Sphingopyxis* as an example for functional validation. In total, we identified 4 ASVs i.e. ASV407, ASV723, ASV3382 and ASV5542 that belong to *Sphingopyxis* at both field locations. We further integrated the results of eGWAS, GWAS, and TWAS to construct a comprehensive multi-omics eQTL regulatory network for relative abundance of *Sphingopyxis*. The network included 416 GWAS significantly associated QTLs, 290 TWAS significant genes of abundance of *Sphingopyxis*, and 697 eQTL hotspots, indicating the complex assembly process of colonization in *Sphingopyxis* rapeseed (Supplemental Dataset 9). To this end, our integrative analyses resulted in nitrogen-related metabolism interactions of host and rhizosphere microbiota. For example, the association detected for ASV723 and variates in the most significantly enriched GO terms “organonitrogen compound metabolic process” (GO:1901564, Adjusted *P* = 0.00652) and “nitrogen compound metabolic process” (GO:0006807, Adjusted *P* = 0.00766) (Supplemental Fig. 11), suggesting a contribution of this taxon to nitrogen homeostasis by modulation of specific metabolism in rapeseed.

To evaluate the function of specific taxon in rapeseed microbiota, we performed a high-throughput bacterial cultivation (Zhang et al., 2021) of the root microbiome from rapeseed varieties YL29 and YL260grown in soil pots. In total, 203 CFUs were recovered and comprised 11 unique bacteria with distinct 16S rRNA gene sequences (Supplemental Dataset 10). These root-derived CFUs represented 4 bacterial phyla and 32 bacterial families associated with rapeseed roots (Fig. 3a). We next mapped the 16S sequence of our cultivated bacteria with the ASVs belonging to *Sphingopyxis* of the large field experiments and only the shared similarity >97% 16S rRNA gene of each ASV was used in the soil inoculation experiment. In particular, for bacterial isolate 29−6000−31 belonging to *Sphingopyxis* the whole genome was assembled with a length of 4,635,643 bp (Fig. 3b). Our comparative COG analyses of the *Sphingopyxis* genome identified a total of 3498 protein-coding genes involved into 222 robust biological pathways enriched in amino acid transport and metabolism category (Supplemental Dataset 11). Finally, we performed single inoculation with *Sphingopyxis* isolate (29−6000−31) derived from rapeseed root in the soil pots under low and high nitrogen levels. Notably, we found that our isolate significantly (*p* = 1.0 × 10^−8^, ANOVA, Tukey HSD) promoted lateral root density of two cultivars (YL29 and YL260) under both nitrogen conditions (Fig. 3c, d). Interestingly, we applied another *Sphingopyxis* isolate (M53) recovered from *Medicago* and this isolate can also partially show the promotion effect on the increase of lateral root density (Fig. 3c, d). We further profiled the metabolome features by targeted metabolomics and identified that inoculation of *Sphingopyxis* reshaped the tryptophan metabolism pathway (Fig. 3e). Interestingly, DR5::GUS imaging demonstrated that inoculation of *Sphingopyxis* modulated auxin transport from the root tip (Supplemental Fig. 12) to the lateral root (Fig. 3f). Importantly, *Sphingopyxis* inoculation showed an influential effect on shoot growth (Fig. 3g) and nitrogen accumulation (Fig. 3h) under low nitrogen conditions. These data suggest that multi-omics is an efficient approach to identify functional bacteria that may contribute to root development and nutrient uptake in rapeseed.

**Figure 3.**
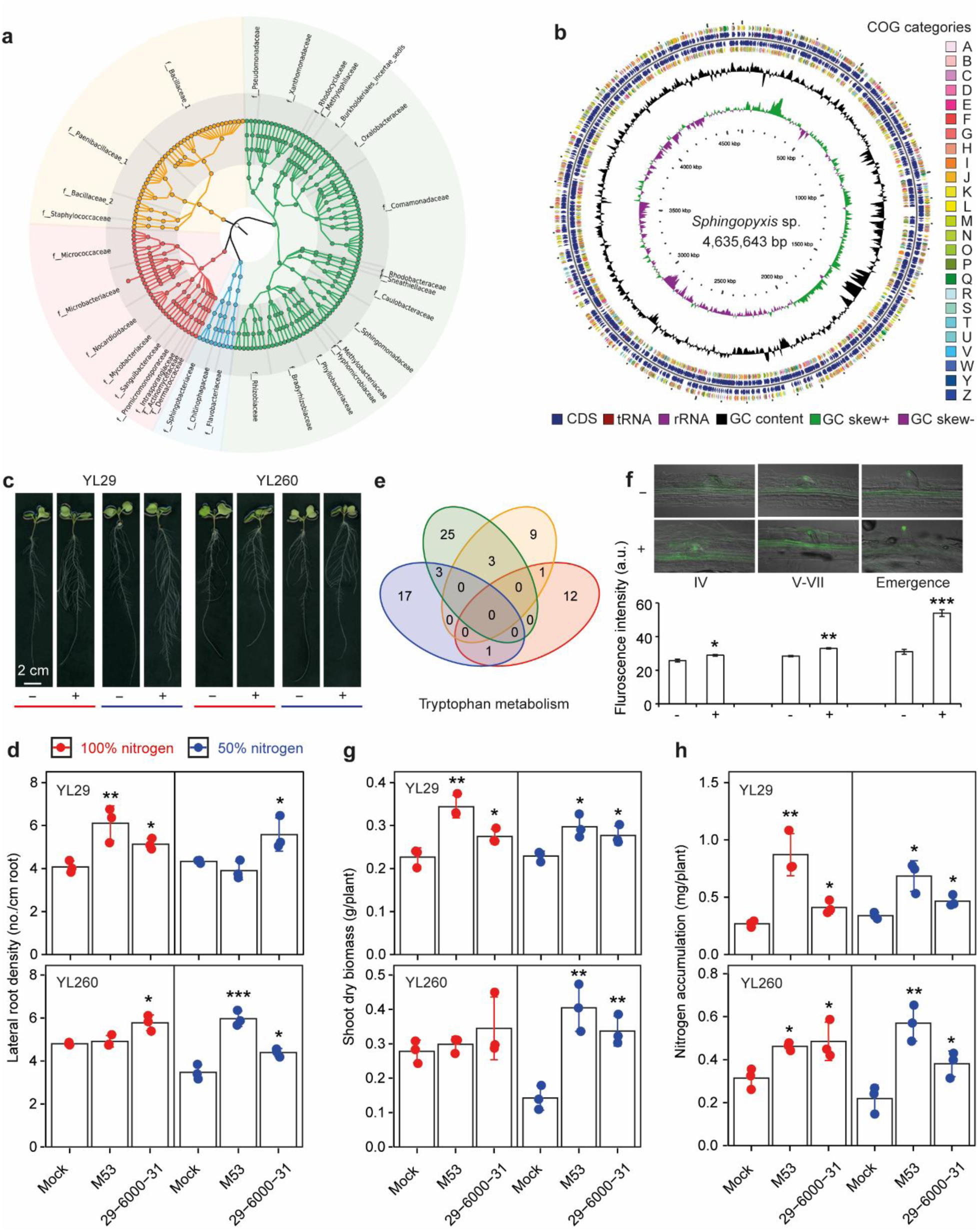
Identification and validation of a novel bacterial genus *Sphingopyxis* which contributes to root development and nitrogen nutrition in *B. napus*. **a,** High-throughput bacterial cultivation and identification from the *B. napus* root system. **b**, Whole genome assembly of the novel bacterial genus *Sphingopyxis*. **c**, Growth performance of *B. napus* genotypes YL29 and YL260 under high (100%) and low (50%) nitrogen conditions with or without *Sphingopyxis* inoculation (isolate 29-6000-31) in the agar system. **d**, Lateral root density of *B. napus* genotypes YL29 and YL260 under high (100%) and low (50%) nitrogen conditions with or without *Sphingopyxis* inoculation (isolate 29-6000-31) in the soil pot system. Here, another bacterial isolate M53 belong to *Sphingopyxis* was derived from *Medicago* root. **e**, Untargeted metabolite profiling identified a core metabolic pathway involved in tryptophan metabolism after *Sphingopyxis* inoculation (isolate 29-6000-31). **f**, DR5::GUS staining of *Arabidopsis* primary root after inoculation with *Sphingopyxis* (isolate 29-6000-31). The lateral root primordium was classified based on different developmental stages. The significances were controlled by paired student *t* tests. *, *P* <0.05; **, *P* <0.01; ***, *P* <0.001. Growth promotion effect on the shoot dry biomass (**g**) and shoot nitrogen content (**h**) after inoculation with *Sphingopyxis* isolates in the soil pot system. The significances were controlled by paired student *t* tests. *, *P* <0.05; **, *P* <0.01; ***, *P* <0.001.

## Discussion

Plants have evolved specialized a rhizosphere dedicated to nutrient acquisition, while harboring a suite of microbes affected by the host’s metabolic properties (Hacquard et al., 2015). Such a reciprocal relationship is genetically controlled in different plant species (Horton et al. 2014; Cordovez et al., 2019; Deng et al., 2021; Escudero-Martinez et al., 2022; Meier et al., 2022; Oyserman et al., 2022; Wang et al., 2022) and heritable (Walters et al., 2018; He et al., 2023), yet the gene regulatory mechanisms remain unknown in plants. This knowledge gap exists for two reasons. First, it is yet unknown to which degree plants can control their gene regulation to guide decisions on microbial selection. For example, legumes are able to form a symbiotic relationship with rhizobia via nod gene transcription (Oldroyd et al., 2011). Yet, whether non-legume crops can develop stable strategies to associate with functional consortia remains obscure. Second, we lack comprehensive studies exploring the effect of the environment in driving the composition and function of the host-associated microbiome, especially in crops. The main reason is that the limited power of small sample size often investigated without considering large population under diverse environments.

We herein present a comprehensive multi-omics analyses of the genetic basis and regulatory landscape of the genome underlying complex plant nutritional traits in association with the rhizosphere microbiome in the crop *Brassica napus* in the field. We adhered to rigorous standards by including a large number of pair-wise datasets (1,341 host SNP array, root transcriptome, rhizosphere 16S rRNA microbiome) and integrating it with 12 root ionome traits under two different environments. Our integrative network association and covariation analyses of host genome, transcriptome, and inhabited microbiome identified causal molecular components and pathways in association with heritable microbial traits linking with root nitrogen status (Fig. 1). We highlight here that plants may contribute to preferential selection of the microbe driven nitrogen nutrition for host benefits among all minerals, thus reflecting its importance in highest abundant element in the atmosphere, and highest requirement for plant lifecycle. We discovered significant genome-wide significant between rhizosphere microbial characteristics and root nitrogen-related genes, in addition to a large number of other host genetic factors, and eventually quantified the total contribution of host genetic loci to microbial variation at each chromosome. Notably, we characterized potential *local*- and *distant*-regulatory variants of gene expression by eQTL mapping, which provides a rich resource to identify genetic and genomic regulatory events involved in host-microbe association (Fig. 2).

In particular, we successfully identified a novel bacterial taxon *Sphingopyxis*, whose variation of abundance trait is largely associated with plant gene expression regulated by eQTL hotspots (Fig. 3). High-throughput cultivation and whole genome assembly provided the full genomic information of *Sphingopyxis*, which displayed a wide range of functional characteristics such as nitrogen and hormone metabolism. Targeted metabolite profiling and DR5::GUS imaging assay demonstrated that *Sphingopyxis* manipulates plant auxin levels to balance lateral root development, which is consistent with the findings of complex hormonal regulation of beneficial root-microbiota interactions in *Arabidopsis* (Finkel et al., 2020; Conway et al., 2022; Gonin et al., 2023). We finally demonstrated that specific *Sphingopyxis* strains facilitate root development and nitrogen acquisition from the bacterial isolates derived from rapeseed and *Medicago* under different conditions (Fig. 3). Indeed, root development and differentiation could coordinate microbiota assembly and plant ionomic homeostasis (Salas-González et al., 2020). Such specific microbe-driven root development will provide a great growth advantage for crop plants under resource-limited conditions (Zhang et al., 2019; Yu et al., 2021). Our results confirmed that host genetic variation impacts root development and nitrogen uptake and that these advantages will contribute to host performance. These findings will help to close the knowledge gap between host plant and biotic interactions with the rhizosphere, and such functional interactions will provide biological causality to further translate host-microbe association into crop resilience.

## Materials and Methods

### Plant materials and field preparation

A panel of 300 *B. napus* accessions covering global diversity were used in this study (Wu et al. 2019). This panel consists of 187 winter, 55 semi-winter, and 58 spring ecotypes based on genetic information and field observations (Supplemental Fig. 1a). Sample size and ecotype information for each geographical origin are provided in Supplemental Dataset 1. In September 2019, all accessions were sown in a randomized complete block design, with three biological replications, in two field locations: China Kaifeng (KF) (34°47’ N, 114°18’ E) and Yangling (YL) (34°17′ N, 108°03′ E) (Supplemental Fig. 1b). The cropping system in the KF plot is an annual winter rapeseed–summer maize rotation, which is representative of the typical cropping system on the North China Plain. The YL plot is representative for an arid agriculture system. Prior to the experiments, the basic physical and chemical properties of the soil from both plots were determined for the top 30 cm soil (Supplementary Table 1). To facilitate seed germination, both plots were sufficiently irrigated to obtain a suitable soil moisture at least 1 month before sowing. The plant material from the previous season were removed from the fields. The same fertilization was applied and mixed with surface soil by rotary tillage in both plots before sowing as: 75 kg P ha^−1^ (as calcium superphosphate), 75 kg N ha^−1^ (as urea) and 75 kg K ha^−1^ (as potassium sulfate). The planting density was synchronized with approximately 165,500 plants ha^−1^, with a row width of 40 cm. No herbicides or additional water were applied during the growth period.

### Rhizosphere harvest at two locations in the field

During to different climate and soil conditions, in total 175 genotypes were recorded and harvested at both fields with comparable growth and performance, respectively. The rhizosphere samples were harvested from 150-day-old rapeseed plants at the flowering stage at both fields. In detail, whole root systems were carefully dug out from the soil profile with the depth ranging from 30 cm to 60 cm, which depends on the distribution of the main roots in the soil profile. The soil closely attached with the roots was defined as the rhizosphere after rigorously shaking off the loosely attached soil particles. Since rapeseed plants have typically one main root with highly branched lateral roots, we specifically dissected lateral roots to have comparable results for accurate comparison among all genotypes. The tightly adhering soil was carefully detached using a brush without damaging the roots, and defined as the rhizosphere (Lee et al., 2011). Rhizosphere samples were collected from two representative plants randomly distributed in each plot and these two samples were pooled together as a whole. These fresh rhizosphere samples were then sieved through a 1 mm sieve to remove visible root fragments and stored at −80 °C for subsequent DNA extraction. To ensure the high-quality of root RNA samples from the field, the same sampling steps were repeated for another two plants and all soil attached to the roots was immediately washed away. Lateral roots were separated by sterilized forceps and be collected together as described during rhizosphere harvest. The washed lateral roots were then rinsed in sterilized water twice to remove potential contaminations, then dried with clean tissue and frozen in 15 ml facon tubes in liquid nitrogen and stored at –80 °C before RNA extraction. Bulk soil samples across each of the plots were taken in the middle between rapeseed rows at a depth of 30 cm as the control. The bulk soils were fully mixed and a representative proportion was collected for downstream analysis.

### Shoot harvest and ionome profiling

The complete aboveground parts (stem and leaves) of rapeseed plants were harvested for all genotypes on the day of harvest at both fields. Then these parts were heat treated at 105 °C for 30 min, dried at 70 °C to constant weight, weighed as the shoot dry biomass. The heat dried parts were then first roughly grinded in a big laboratory crusher machine (BF-10, Hebei Beichen Technology Co., Ltd), followed by fine grinding to powder using a ball milling machine (Multifunctional ball mill QM100S, Beijing Wuzhou Dingchuang Technology Co., Ltd). In total, concentrations of 12 mineral nutrients were determined as previously described (Yu et al., 2021). In brief, 2−5 mg of fine plant powder was used to profile the total nitrogen concentration in an elemental analyzer (Isotope ratio mass spectrometer, Vario PYRO Cube-IsoPrime 100, Elementar-Isoprime Ltd.). Data were then calculated into peak areas by the software Callidus, providing quantitative results using reference material as a calibration standard. The same plant material was used to determine the concentrations of 11 additional mineral nutrients using the Inductively Coupled Plasma Optical Emission Spectrometer (ICP-OES) machine (19A07591, Agilent Technology Co. Ltd).

### Genomic DNA extraction and bacterial community profiling in the rhizosphere

Genomic DNA was extracted from a total of 1,050 rhizosphere samples derived from 175 genotypes at two field locations using the E.Z.N.A.^®^ soil DNA Kit (Omega Bio-tek, Norcross, GA, U.S.) according to the manufacturer’s protocol. All DNA samples were quality checked and the concentration was quantified by NanoDrop 2000 spectrophotometers (Thermo Fisher Scientific, Wilmington, DE, USA). Bacterial 16S rRNA gene fragments (the hypervariable region V3−V4) were amplified using primers 338F (5’-ACTCCTACGGGAGGCAGCAG-3’) and 806R (5’-GGACTACHVGGGTWTCTAAT-3’) at the following PCR conditions: an initial denaturation at 95 °C for 30 s, followed at 95 °C for 30 s, 55 °C for 30 s and 72 °C for 45 s for 27 cycles, then single extension at 72 °C for 45 s and storage of samples at 4 °C . PCR was performed with 4 μL 5 × TransStart FastPfu buffer, 2 μL 2.5 mM deoxynucleoside triphosphates (dNTPs), 0.8 μL of each oligonucleotide primer (5 μM), 0.4 μL TransStart FastPfu DNA Polymerase, 10 ng of extracted DNA, and finally ddH_2_O to obtain a volume of 20 μL. Agarose gel electrophoresis was performed to verify the size of amplicons. Amplicons were subjected to paired-end sequencing on the Illumina MiSeq PE300 platform.

After demultiplexing, the resulted sequences were merged with FLASH (v1.2.11) (Magoč and Salzberg 2011) and quality filtered with fastp (v0.19.6) (Chen et al. 2018). The qualified sequences were de-noised using the DADA2 (Callahan et al. 2016) plugin in the Qiime2 (v2020.2) (Bolyen et al. 2019) pipeline with default parameters, which obtains single nucleotide resolution based on error profiles within samples. Denoised sequences are usually called amplicon sequence variants (ASVs). Taxonomic assignment of ASVs was performed using the sklearn-based Naive Bayes classifer implemented in Qiime2 and the SILVA 16S rRNA database (v138). Based on the ASVs information, rarefaction curves and alpha diversity indice Shannon index was calculated with Mothur (v1.30.1). The similarity among the microbial communities in different samples was determined by principal coordinate analysis (PCoA) based on Bray-Curtis dissimilarity using R Vegan (v2.4-3) package. The PERMANOVA test was used to assess the percentage of variation explained by the treatment along with its statistical significance using R Vegan (v2.4-3) package.

### Root RNA sequencing and gene expression analysis

Total RNA was extracted from the root tissue using TRIzol® reagent according the manufacturer’s instructions (Invitrogen) and genomic DNA was removed using DNase I (TaKara). Then RNA quality was determined by 2100 Bioanalyser (Agilent). Only high-quality RNA sample (RIN> 8.0) was used for library construction. RNA-seq library was prepared following TruSeqTM RNA sample preparation Kit from Illumina (San Diego, CA) using 1μg of total RNA. Briefly, mRNA was isolated according to the polyA selection method by oligo(dT) beads and then fragmented by fragmentation buffer. Subsequently, double-stranded cDNA was synthesized using a SuperScript double-stranded cDNA synthesis kit (Invitrogen, CA) with random hexamer primers (Illumina). Then the synthesized cDNA was subjected to end-repair, phosphorylation and ‘A’ base addition according to Illumina’s library construction protocol. Libraries were selected for cDNA target fragments of 300 bp on 2% Low Range Ultra Agarose followed by PCR amplified using Phusion DNA polymerase (NEB) for 15 PCR cycles. After quantified by TBS380, paired-end RNA-seq sequencing libraries were sequenced with the Illumina NovaSeq 6000 sequencer (2 × 150bp read length).

The raw paired end reads were trimmed and quality controlled by fastp (Version 0.19.5, https://github.com/OpenGene/fastp) with default parameters (Chen et al. 2018). Then clean reads were separately aligned to the reference genome (Brassica napus: https://www.genoscope.cns.fr/brassicanapus/data/) with orientation mode using HISAT2 (Version 2.1.0, http://ccb.jhu.edu/software/hisat2/index.shtml) software (Kim et al. 2015). The expression level of each transcript was calculated according to the transcripts per million reads (TPM) method. RSEM (Version 1.3.1, http://deweylab.biostat.wisc.edu/rsem/) was used to quantify gene expression (Li and Dewey 2011).

### WGCNA and correlation with phenotypic traits

To better understand the causal relationships between different microbial assemblies and taxonomic shifts among samples, we applied an unbiased data-driven method i.e. weighted correlation network analysis (WGCNA) to explore the biological relevance between gene expression and microbial traits (Langfelder and Horvath, 2008). We subjected WGCNA (1.72.1) in R to identify different gene expression modules, which represented different patterns of correlation across location and genotypes within the root. We filtered and normalized the gene table to construct a robust network with co-expressed gene modules. The soft thresholding power β was determined by default and used to calculate adjacency matrix. Moreover, we transformed the adjacency matrix into a topological overlap matrix (TOM) with selected power and calculated the corresponding dissimilarities (dissTOM) as “1 – TOM” to minimize the effects of noise. We then used dissTOM as a distance measure and set the minimum module size (number of ASVs to 30) to detect gene co-expressed modules. In addition, we quantified the similarities of entire modules and calculated the “eigengene”, which were subsequently used to associate with different ionomic traits and microbial ASVs. We chose modules that have a Spearman correlation coefficient >0.1 and *P* value <0.05 with different traits as significant associated modules. Network visualization was performed in Cytoscape (3.8.0) only for significant modules.

### Heritability estimation for microbial ASVs

We calculated the broad-sense heritability of all ASVs that were subjected to samples with at least 10,000 reads across at least 80% of samples. The relative abundances of ASVs were normalized and log-transformed to fit the normal distribution. The following model was used to estimate the heritability based on He et al., 2023:

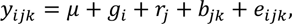

where 𝑦_𝑖𝑗𝑘_ is the observation of the *i*-th genotype in the *k*-th block of the *j*-th complete replicate. 𝜇 is the general mean, 𝑔_𝑖_ is the effect of the *i*-th genotype, 𝑟_𝑗_ is the effect of the *j*-th replicate, 𝑏_𝑗𝑘_ is the effect of the *k*-th block nested within the *j*-th replicate and 𝑒_𝑖𝑗𝑘_ is the residual term. All effects except the general mean were assumed to be random and follow an independent normal distribution.

The heritability was calculated using the following formula:

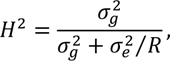

where 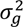 and 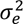 are the estimated genotypic and residual variance, *R* is the number of replications. The best linear unbiased estimations (BLUEs) of all genotypes for each ASV at each location were obtained by fitting model once more, assuming the general mean and genotypic effects are fixed and all other effects are random. All linear mixed models were fitted using the software ASReml-R 4.0 (Butler et al., 2017).

### Linkage disequilibrium

Linkage disequilibrium (LD) decay with physical distance between pairwise SNPs was calculated in a 500 kb window and visualized by using PopLDdecay software (Zhang et al., 2019b) with default parameters for the whole genome sequencing (WGS) SNPs and transcriptome sequencing (RNAseq) SNPs in KF and YL, respectively. LD decay to 0.1 report as the measure of decay, at around of 1.5 kb and 50 kb for WGS and RNAseq SNPs data (Supplementary Figure 13). For convenience of claim the QTL region, 50 kb was used in the GWAS and eQTL analysis.

### GWAS on ASVs

To test the association between genomic regions and rhizosphere microbiomes, the abundance of microbial ASVs were treated as quantitative trait. And, for the genotypic data, after imputation by Beagle v5.2 (Browning et al., 2018), in total, 345,289 WGS SNPs with minor allele frequency of ≥0.05 and a genotype heterozygosity rate of ≤95% in the entire population were used for GWAS. A standard single variant (SNP)-based “Q+K” GWAS analysis were performed by a mixed linear model (Yu et al., 2006) for all 203 ASVs. A marker-derived kinship matrix generated was used to correct the structure of multiple levels of relatedness, or control the polygenic background effects. More precisely, the model is of the following form:

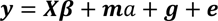

where 𝒚 is the *n*-dimensional vector of phenotype data of ASV (i.e. BLUEs of ASV within a certain environment (such as KF or YL), *n* is the number of genotypes), 𝜷 is the *k*-dimensional vector of fixed covariates including the common intercept and 𝑿 is the corresponding *n* × *k* design matrix. 𝑎 is the additive effect of the marker being tested, 𝒎 is the *n*-dimensional vector of marker profiles for all genotypes. The elements in 𝒎 are coded as 0, 1 or 2, which is the number of minor alleles at the SNP. 𝒈 is an *n*-dimensional random vector representing the genetic background effects. We assume that 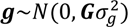, where 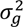 is the genetic variance component, 𝑮 is the VanRaden genomic relationship matrix (VanRaden et al., 2008). 𝒆 is the residual term and 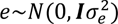, where 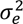 is the residual variance component and 𝑰 is the *n* × *n* identity matrix. After solving the linear mixed model, the marker effect was tested using the Wald test statistic 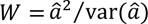, which approximately follows a 𝜒^2^ -distribution with one degree of freedom. To reduce the computational load, we implemented a commonly used approximate approach, namely the “population parameters previously determined” (P3D) method (Zhang et al., 2010). That is, we only fit the model once without any marker effect, and then we fixed the estimated the variance parameters 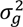 and 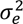 throughout the testing procedure. Then, the test statistic for each marker can be efficiently calculated. All analysis procedure was implemented using R codes developed by ourselves. The variance parameters were estimated by the Bayesian method using the package BGLR (Pérez and de Los Campos 2014).

For each microbial ASVs, the significant marker-trait association was identified with a threshold of *p*-value < 0.05 after FDR (false discovery rate) correction for multiple test (Benjamini and Hochberg, 1995). The *p*-value after FDR correction was used for Manhattan plot, while the original *p*-value for QQ-plot.

Considering the Linkage disequilibrium (LD) between significant SNPs and/or its neighboring SNPs, the significant SNPs need to be traced back and grouped into independent QTL regions. For this, these steps were followed to identify QTL interval. First, the upstream and downstream LD boundaries of each significant SNP were identified based on the LD decay between the significant SNPs and its neighboring SNPs, for instance, the genomic position where the LD first dropped below 0.1. Then, each significant SNP has its own QTL boundaries. Next, those candidate QTL region with overlap or the distance between peak SNPs less than 50 kb (the average distance of LD decay to 0.1 is about 50 kb) were merged into independent QTL, represented by their most significant SNP (named as lead SNP).

### The identification of local eQTL and distant eQTL hotspots

One of the assumptions of detecting the association between SNPs and gene expression through linear mixed model is that the residual of expression values follows a normal distribution in each genotype class, which is violated by outliers or non-normality in gene expression estimated from the sequencing reads. For each gene, the expression was normalized by QQ-normal of R package. Finally, both in KF and YL, a dataset including 17006 genes were obtained and the gene expression level would be treated as expression traits for eGWAS analyses. For the SNPs data, after imputation by Beagle v5.2 (Browning et al., 2018), in total, 239,172 RNAseq SNPs (or 292,839 RNAseq SNPs) with minor allele frequency of 0.05 or greater and a genotype heterozygosity rate of 95% or less in the entire population were used for eGWAS in KF (or YL). The SNP-gene expression association test statistic was performed by a standard “Q+K” mixed linear model (Yu et al., 2006) same as descripted elsewhere above. And, a SNPs-derived VanRaden genomic kinship matrix (VanRaden et al., 2008) also generated was used to correct the structure of multiple levels of relatedness, or control the polygenic background effects. For each gene expression traits, the significant association between a SNP (termed an eSNP) and the expression of a gene (termed an eGene) was identified with a threshold of *p*-value < 0.05 after Bonferroni-Holm correction for multiple test (Holm, 1979). More precisely, the number of tests equaled the number of markers, i.e., 239,172 for KF and 292,839 for YL, and the corresponding threshold was 2.09×10^-7^ and 1.71×10^-7^, respectively. The same approach (trace SNP to QTL as descripted above) was used to trace back the eSNP to eQTL, the genomic region significantly affects the expression level of eGene.

To classify the eQTL as local eQTL and distant eQTL, the physical genomic region of the eQTL and their target genes were compared. If the start of eQTL located within 50 kb downstream of the eGene, or the end of eQTL located within 50 kb upstream of the eGene, it was considered a local eQTL, otherwise a distant eQTL.

To identify the potential distant eQTL hotspot regions at KF and YL, hot_scan software (Silva et al., 2014) was used for all eQTL genomic region in each chromosome with the following parameters: -m 100000 -s 0.01. The distant eQTL hotspots were visualized using R package circlize v0.4.13 (Gu et al., 2014).

### Transcriptome-wide association study for ASVs

Transcriptome-wide association studies (TWAS) integrate genome-wide association studies (GWAS) and gene expression datasets to identify gene–trait associations (Gusev et al., 2016). To test the association between gene expression level and rhizosphere microbiomes, the abundance of microbial ASVs were treated as quantitative trait and the expression level of 17006 genes at KF and YL were treated as explanatory variants. The association analysis was also performed by a standard “Q+K” mixed linear model (Yu et al., 2006) same as described above. However, to account for hidden batch effects or other confounders in the expression data and control the structure of multiple levels of relatedness, we utilized a gene expression-derived kinship matrix. Specifically, the kinship matrix was calculated by 𝑲_𝑒𝑥𝑝_ = 𝑀𝑀^′^/𝑐, and 𝑀 is the matrix of standardized features of gene expression, 𝑐 is the mean of all diagonal elements in the matrix 𝑀𝑀^′^.

Following the same approach to solve the mixed linear model, the *P* values were calculated for all genes for each ASV. Subsequently, the raw *P* values were adjusted to control the false discovery rate (FDR) using the p.adjust() function in base R v4.1.0 (Benjamini and Hochberg, 1995; R Core Team, 2018). Significant associations were identified as those passing a 5% FDR threshold for TWAS.

Both for GWAS, eGWAS and TWAS, quantile–quantile plots all showed that the genomic background was properly controlled in the mixed linear model, thereby avoiding inflated rates of false-positive QTL, eQTL and gene–ASV associations.

### Construction of the ASV-GWAS-eGWAS-TWAS regulated network

Based on the QTL, hotspot results and TWAS significant genes, a genomic regulating ASV network was constructed. The connection between the nodes including QTL, Hotspot, Gene and ASV were visualized using Gephi v0.10 ( https://github.com/gephi/gephi). And then, we constructed the Bn-ASV-QTL-eQTL-gene database (Supplementary Datasets 7-10).

### Prediction for ASVs and ionomic traits using multiple omics data

To explored the possibility of predicting the ionome traits in each location (KF and YL), four omic (genomic data SNPs calling from whole genome sequencing data and RNA sequencing data, transcriptomic and microbiome ASVs) data collected from 175 *Brassica napus* accessions were used for prediction. All accessions were genotyped using WGS (Wu et al., 2019). RNA sequencing was subsequently performed for these 175 lines using 150 bp pair-end Illumina reads at each location. A total of 752,414 SNPs from WGS, 243,209 SNPs from RNA-seq in KF and 295,925 SNPs from RNA-seq in YL were obtained. Moreover, 52,962 gene expression features (transcriptomic data) were obtained. Microbiome ASVs profiling was carried out in the rhizosphere of rapeseeds and 2,319 highly abundant and heritable ASVs were detected at each location.

After removing the SNPs without polymorphism and Beagle v5.2 (Browning et al., 2018) imputation, we obtained 637,823 SNPs from WGS, 241,558 SNPs from RNA-seq in KF and 295,925 SNPs from RNA-seq in YL. 17,006 genes and 203 ASVs were selected for subsequent prediction analysis both in KF and YL. We analyzed 13 ionome traits (Al, B, Ca, Cu, Fe, K, Mg, Mn, Na, P, Zn, N, and C) to evaluate the prediction ability.

In Both KF and YL, using four types of omics data and considering all possible combination, we have 15 cases, in which the ionome phenotypes were predicted using one types of data, WGS SNPs (WGS), RNA SNPs (RNA), gene expressions (Gene), and microbiome data (ASV); using two types of data, WGS+RNA, WGS+Gene, WGS+ASV, RNA+Gene, RNA+ASV, and Gene+ASV; using three types of data, WGS+RNA+Gene, WGS+RNA+ASV, WGS+Gene+ASV, and RNA+Gene+ASV; and using all four types data, WGS+RNA+Gene+ASV. The model can be uniformly described as follows within each location:

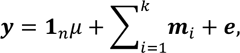

where 𝒚 is the *n*-dimensional vector of ionome phenotypic records (i.e. BLUEs within a certain location, n is the number of genotypes), 𝒎_𝑖_ is an n-dimensional trait values for all individuals determined by a certain combined type of omics data in a specific sample, *k* can be 1 (WGS, RNA, Gene, or ASV), 2 (WGS_RNA, WGS_Gene, WGS_ASV, RNA_Gene, RNA_ASV, or Gene_ASV), 3 (WGS_RNA_Gene, WGS_RNA_ASV, WGS_Gene_ASV, or RNA_Gene_ASV) or 4 (WGS_RNA_Gene_ASV). ***e*** is the residual term and 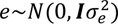, where 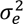 is the residual variance component and ***I*** is the n × n identity matrix. We assume that 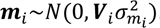, where 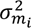 is the corresponding variance component, 𝑽*_i_* is the VanRaden genomic relationship matrix (VanRaden et al., 2008) for WGS and RNA SNPs data. For gene expression and ASV data, 𝑽_𝑖_ is a covariance matrix derived from the corresponding omics data. Assuming that 𝑴_𝑖_ is the *n* × *t* matrix of standardized features of gene expression or ASVs (t is the number of features), we have 𝑽_𝑖_ = 𝑀_𝑖_𝑀_𝑖_^′^/𝑐_𝑖_ where 𝑐_𝑖_ is the mean of all diagonal elements in the matrix 𝑴_𝑖_𝑴_𝑖_^′^.

The prediction ability was evaluated in a five-fold cross validation scenario. Prediction models were implemented using the R package BGLR (Pérez and de Los Campos 2014). Cross validation was repeated 100 times and the prediction ability was estimated as the Pearson correlation coefficient between the observed and the predicted ionome phenotypic values. Meanwhile, a similar strategy was used to predict the abundance of ASVs. With the input of three type of omics data (WGS SNPs (WGS), RNA SNPs (RNA), gene expressions (Gene)), we have 7 combinations of input data to predict the ASV, namely, WGS, RNA, Gene, WGS_RNA, WGS_Gene, RNA_Gene and WGS_RNA_Gene. A similar model and five-fold cross validation were used to evaluate the prediction ability.

### Transcriptome, microbiome and ionome association analysis

To estimate the correlation between root gene expression, ionome accumulation and rhizosphere microbiome assembly, we performed the Mantel statistical test among these data matrices. Euclidean distance was calculated for root gene expression and ionome accumulation, and Bray-Curtis distance was determined for microbial composition. The Pearson correlation method was used in ‘mantel’ function in vegan package (v2.6.4) of R. Permutations = 1999.

### Functional characterization of genes associated with *Sphingopyxis*

To understand the function of genes associated with the genus *Sphingopyxis*, we performed gene ontology (GO) enrichment analyses with the genes identified by GWAS and TWAS analyses. GO enriched terms were identified using Goatools (https://github.com/tanghaibao/goatools) and showed significant enrichment in GO terms with an FDR <0.05 compared to the whole-transcriptome background.

### Whole genome assembly and annotation for *Sphingopyxis*

To deepen the understanding of the genetic and genomic information of *Sphingopyxis*, we *de novo* assembled the whole genome of bacterial isolates derived from *Sphingopyxis* using Nanopore and Illumina platforms. Bioinformatic analyses were performed using the online Majorbio Cloud (http://cloud.majorbio.com, Ren et al., 2022) from Shanghai Majorbio Bio-pharm Technology Co.,Ltd. The detailed procedures are as described as following. The raw Illumina sequencing reads generated from the paired-end library were subjected to quality-filtered using fastp (v0.23.0). Nanopore reads were extracted, basecalled and demultiplexed, and trimmed using ONT Guppy with the minimum Q score cutoff of 7. Then the clean short and long reads were co-assembled to construct complete genomes using Unicycle (v0.4.8) (Wick et al., 2017). As a final step, Unicycler uses Pilon (v1.22) was used to polish the assembly using short-read alignments, reducing the rate of small errors. The coding sequences (CDs) of chromosome and plasmid were predicted using Prodigal (v2.6.3) (Hyatt et al., 2010) respectively. tRNA-scan-SE (v 2.0) (Chan et al., 2021) was used for tRNA prediction and Barrnap (v0.9) (https://github.com/tseemann/barrnap) was used for rRNA prediction. The predicted CDs were annotated from COG (https://www.ncbi.nlm.nih.gov/research/COG) and KEGG database (https://www.genome.jp/kegg/kegg1.html) using sequence alignment tools Diamond (Buchfink et al., 2021) and HMMER (Finn et al., 2011). Each set of query proteins were aligned with the databases and annotations of best-matched subjects (e-value <10^-5^) were obtained for gene annotation.

### High-throughput bacterial cultivation and identification by Illumina sequencing

To obtain representative bacteria for functional validation, we performed high-throughput bacterial cultivation (Zhang et al., 2021) using two representative rapeseed varieties i.e. YL29 and YL264 that were grown in natural soil at the experimental station at Yangling, where one of the field experiments was carried out. These two genotypes were harvested at flowering stage to synchronize with our field sampling. In detail, the fresh dug out roots of three representative plants of each variety were mixed thoroughly to represent both varieties and reduce sample variation. The fully washed roots were further rinsed three times in sterile washing buffers (PBS) on a shaker at 180 rpm. for 15 min. Then the root tissue was ground into homogeneous slurry, which was transferred into MgCl_2_ to sediment for 15 min. The supernatants were diluted, distributed and cultivated in 96-well cell culture plates in 1:10 (v/v) tryptic soy broth for 15 days at room temperature. The dilution gradients were determined by preliminary experiments and appropriate concentration of diluent (6000×) was retained for subsequent bacterial identification for the genotype YL29 and YL264.

To identify pure bacterial cultures, we next carried out a two-sided barcoded PCR combined with Illumina MiSeq for sequencing the bacterial 16S rRNA (V5–V7) gene. DNA of the cultivated bacteria was extracted using the alkaline lysis method (Zhang et al., 2021). In detail, 6 μl of bacterial cultures was added to 10 μl lysis buffer I containing 25 mM NaOH and 0.2 mM Na_2_-EDTA, at pH 12 at 95 °C for 30 min incubation. Then the pH value was adjusted to 7.5 by addition 10 μl of buffer II containing 40 mM Tris-HCl. Individual positions of each isolate in 96-well plates were indexed by a two-step PCR protocol using the degenerate primers 799F and 1193R containing well- and plate-specific barcodes to amplify the variable regions V5–V7. In the first PCR, degenerate primers were used to amplify the bacterial 16S rRNA gene. We set up the well positions A12 and B12 for the negative and positive controls in each plate of bacterial DNAs by adding nuclease-free water and *E. coli* DNA as amplification templates. The PCR reaction mixture and amplification procedure for each well was performed accordingly. The first PCR products of each 96-well were diluted 40× and used as templates and each PCR product was labelled with barcoded primers in the second PCR. The PCR reaction system and cycling conditions were described previously (Zhang et al., 2021). The second PCR products were purified using the Wizard SV Gel and PCR Clean-up System (Promega) and the Agencourt AMPure XP Kit (Beckman Coulter). DNA concentration was measured by Quant-iT PicoGreen dsDNA Assay Kit (Life Technologies) and samples were pooled in equal amounts. A total of 1500 ng of purified PCR product libraries were sequenced on an Illumina MiSeq platform. Each sequence contained a plate barcode, a well barcode and V5–V7 sequences. Bioinformatics analyses to identify the bacterial taxa was performed according to an established pipeline (Zhang et al., 2021).

### Purification and preservation of cultivated bacteria

To identify specific ASVs from cultivated bacteria, five wells containing the corresponding bacteria were selected and inoculated on a solid media for three consecutive purifications before an individual colony was used for liquid cultures. Then these liquid cultures were used to validate the bacterial 16S rRNA gene sequence by Sanger sequencing with both 27F (AGAGTTTGATCCTGGCTCAG) and 1492R (TACGGCTACCTTGTTACGACTT) primers as well as for the preservation of glycerol stocks for the cultivated bacteria. Phylogenetic relationships of cultivated bacterial isolates were constructed using MEGA11 software.

### Metabolome profiling from bacteria-inoculated rapeseed roots

To explore whether inoculation of *Sphingopyxis* will influence the metabolic feature of root system, we performed an untargeted metabolome approach for rapeseed plants grown in the same natural soil as described above. In detail, the rapeseeds used in the experiment were cultivated under two nitrogen conditions, e.g. half nitrogen (2.375 mM KNO_3_, 11.775 mM KCl, 313 μM KH_2_PO_4_, 750 μM CaCl_2_, 50 μM H_3_BO_3_, 50 nM CuSO_4_•5H_2_O, 50 μM MnSO_4_•H_2_O, 2.5 μM KI, 375 μM MgSO_4_, 52.5 nM CoCI_2_•6H_2_O, 44 μM FeNaEDTA, 0.5 μM Na_2_MoO_4_•2H_2_O, 15 μM ZnSO_4_•7H_2_O) and full nitrogen (4.75 mM KNO_3_, 9.4 mM KCl, 313 μM KH_2_PO_4_, 750 μM CaCl_2_, 50 μM H_3_BO_3_, 50 nM CuSO_4_•5H_2_O, 50 μM MnSO_4_•H_2_O, 2.5 μM KI, 375 μM MgSO_4_, 52.5 nM CoCI_2_•6H_2_O, 44 μM FeNaEDTA, 0.5 μM Na_2_MoO_4_•2H_2_O, 15 μM ZnSO_4_•7H_2_O) supply for genotypes YL29 and YL260 for 28 days. Half of the plants were inoculated with *Sphingopyxis* isolate that originated from the cultivation. Before inoculation, we have confirmed the 16S sequence of the isolates with the generated ASVs from the field experiments using HISAT2 (Kim et al., 2019). The preparation and inoculation were performed according to our previous work (Yu et al., 2021). For sample harvest, soil from the whole root systems were completely washed off and rinsed by sterile water for three times. In particular, we cut off the lateral roots, dried them with clean tissue and immediately froze them in liquid nitrogen. Around 100 mg of lateral roots were ground into fine powder in liquid nitrogen and mixed with prechilled 80% methanol, vortexed, and incubated on ice for 5 min. This was followed by centrifugation at 15,000 *g*, at 4 °C for 20 min. The supernatant was diluted to a final concentration containing 53% methanol by LC-MS grade water and transferred to a fresh Eppendorf tube. After centrifugation at 15,000 *g*, 4 °C for 20 min, the supernatant was injected into the LC-MS/MS system for analysis (Want et al., 2012).

The sample were analyzed by LC-MS/MS using an ExionLC™ AD system (SCIEX) coupled with a QTRAP® 6500+ mass spectrometer (SCIEX) in Novogene Co., Ltd. (Beijing, China). A 20-min linear gradient was used to inject samples onto an Xselect HSS T3 (2.1 × 150 mm, 2.5 μm) at a flow rate of 0.4 ml/min for positive/negative polarity mode. The eluents were eluent A (0.1% Formic acid-water) and eluent B (0.1% Formic acid-acetonitrile) (Ping et al., 2015). The gradient elution technique was employed with the following setting: 2% B, 2 min; 2-100% B, 15 min; 100% B, 17 min; 100-2% B, 17.1 min; 2% B, 20 min. QTRAP® 6500+ mass spectrometer was provided in both negative and positive ionisation modes with Curtain Gasof 35 psi, Collision Gas of Medium, Temperature of 550 °C, Ion Source Gas of 1:60, Ion Source Gas of 2:60, and IonSpray Voltage of 5500V in positive polarity mode and −4500V in negative mode. Based on the in-house database from Novogene, the samples were detected by MRM (Multiple Reaction Monitoring). Quantitative analysis of metabolites was performed by Q3, while qualitative analysis was performed using Q1, Q3, RT (retention time), DP (clustering potential) and CE (collision energy). HPLC-MS/MS generated data files processed with the SCIEX OS Version 1.4 to integrate and correct the peak under the following parameters: minimum peak height, 500; signal/noise ratio, 5; gaussian smooth width, 1. The area of each peak represents the relative content of the corresponding substance. Annotation of metabolites based on KEGG database (http://www.genome.jp/kegg/). Partial least squares discriminant analysis (PLS-DA) were performed at metaX (Wen et al., 2017). The statistical significance was estimated using univariate analysis (Student’s *t* test). Differential metabolites were defined when VIP >1 and P <0.05 and fold change ≥2, or ≤0.5. Metabolic pathway enrichment was performed and considered enriched when ratios were satisfied by x/n >y/N and statistically significant when the *P* value of metabolic pathway <0.05.

### Confocal microscopy for DR5::VENUS expression in the root

*Arabidopsis thaliana* Columbia-0 (Col-0) was used in this study. Arabidopsis seeds were surface-sterilized with 75% (v/v) ethanol for 2 min, washed three times with sterile water, then soaked in 10% (v/v) sodium hypochlorite for 5 min, and washed five times with sterile water. Individual bacterial strains were cultured in Tryptic Soy Broth medium (TSB, Oxoid) for 24 h. Bacterial cells were centrifuged, washed twice and resuspended in sterile water and adjusted to a final density of 108 CFU ml-1 (OD600 = 0.5). Sterilized seeds were sown on plates (13cm × 13cm) containing 50ml 1/4 Murashige & Skoog (MS, Coolaber) medium supplemented with 10 g/L sucrose and 10 g/L agar. After 2 d of stratification, plates were positioned vertically in a growth chamber under a long-day photoperiod (16 h: 8 h, light: dark, relative humidity 60%) at 22 °C.

### Nitrogen phenotype screening in the agar-plate system

The seeds of genotypes YL29 and YL260 were surface-sterilized with 75% (v/v) ethanol for 2 min, washed three times with sterile water, then soaked in 10% (v/v) sodium hypochlorite for 5 min, and washed five times with sterile water. Individual bacterial strains were cultured in Tryptic Soy Broth medium (TSB, Oxoid) for 24 h. Bacterial cells were centrifuged, washed twice and resuspended in sterile water and adjusted to a final density of 10^8^ CFU ml^-1^ (OD_600_ = 0.5). Sterilized seeds were sown on plates (25 cm × 25 cm) containing 200ml 1/4 Murashige & Skoog (MS, Coolaber) medium supplemented with 10 g/L sucrose and 10 g/L agar. We defined the concentration of nitrogen in the all-element 1/4MS medium as 100% nitrogen and 50% nitrogen. In brief, the MS medium was autoclaved at 120 °C for 20 min. 1 ml bacterial suspension or 1ml sterile water as mock was added to the medium when the temperature drops to about 55 °C. The mixture was shaken well and poured it into the plate. The seeds were evenly distributed on the plates, which were sealed with Parafilm and incubated in a growth chamber under a long-day photoperiod (16h: 8h, light: dark, relative humidity 60%) at 25 °C. After 9-day culture, the root traits including the total length, lateral root density and shoot traits including biomass, nitrogen accumulation were investigated according to Yu et al., (2021).

### Functional validation experiments in the soil pots

For the pot experiment, the soil matrix (soil: vermiculite: perlite = 6: 3: 1) was sterilized twice with an interval of 24 h. The seeds of genotypes YL29 and YL260 were surface-sterilized according to the method described above and cultivated on the plate for 24 h until the seeds germinated. Bacterial cells were centrifuged, washed twice and resuspended in phosphate buffer and adjusted to a final density of 10^8^ CFU ml^-1^ (OD_600_ = 0.5). The germinated seeds were immersed in the bacterial suspension for two hours, and the control was immersed in sterile phosphate buffer. After that, four germinated seedlings were planted in each pot containing 200g soil in a growth chamber under a long-day photoperiod (16h: 8h, light: dark, relative humidity 60%) at 25 °C. After transferring, each seedling was watered with 2 ml bacterial suspension, and the control was watered with 2 ml phosphate buffer. After 15 days, 5 ml of nutrient solution with different nitrogen gradients poured into each pot, and the soil was kept moist by daily watering. After 28 days of transplantation, the root and short samples were harvested as described previously.

## Data availability

All raw rapeseed RNAseq and bacterial 16S gene data generated in this paper were deposited in the Sequence Read Archive (http://www.ncbi.nlm.nih.gov/sra) under the BioProject IDs PRJNA986524 (KF, RNAseq), PRJNA990484 (YL, RNAseq), PRJNA956663 (KF, 16S) and PRJNA960662 (YL, 16S).

## Acknowledgements

We thank the Institute of Crop Science of Zhejiang University for providing all *Brassica napus* L. ecotypes. We are grateful for the generous contribution of DR5::GFP *Arabidopsis* marker lines by Nicolaus von Wirén (IPK Gatersleben, Germany). This work was supported by the National Natural Foundation of China (grant numbers: 32072102, 31671728, 32130076), the National Key R & D Program of China (2022YFD1900705), the Chongqing Talent Funding to N. L., the Deutsche Forschungsgemeinschaft (DFG) grant 514003603, the DFG Emmy Noether Programme 444755415, the German Excellence Strategy – EXC 2070 – grant 390732324 to P.Y. and DFG Priority Program (SPP2089) “Rhizosphere Spatiotemporal Organisation - a Key to Rhizosphere Functions” grant HO2249/17-2 to F.H. and P.Y., the Fundamental Research Funds for the Central Universities (SWU-XJLJ202308; XDJK2018AA005), Shuangcheng Cooperative Agreement Research Grant of Yibin, China (XNDX2022020003) and the China Postdoctoral Science Foundation (Grant No. 2020M672213). We thank New Cornerstone Science Foundation through the XPLORER PRIZE to Y.B.

