## Supplemental information for "Large-scale multi-omics analyses identified root-microbiome associations underlying plant nitrogen nutrition"

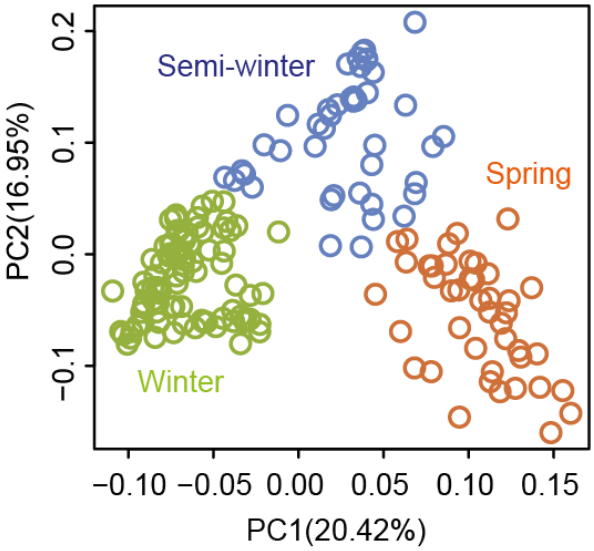


**Supplemental Figure 1. A principal component analysis for 175 *B. napus* ecotypes grown under two field trial locations in China**. The ecotypes are classified as Spring, Winter and Semi-winter. The locations were established in Kaifeng, North China and Yangling, Northwest China.

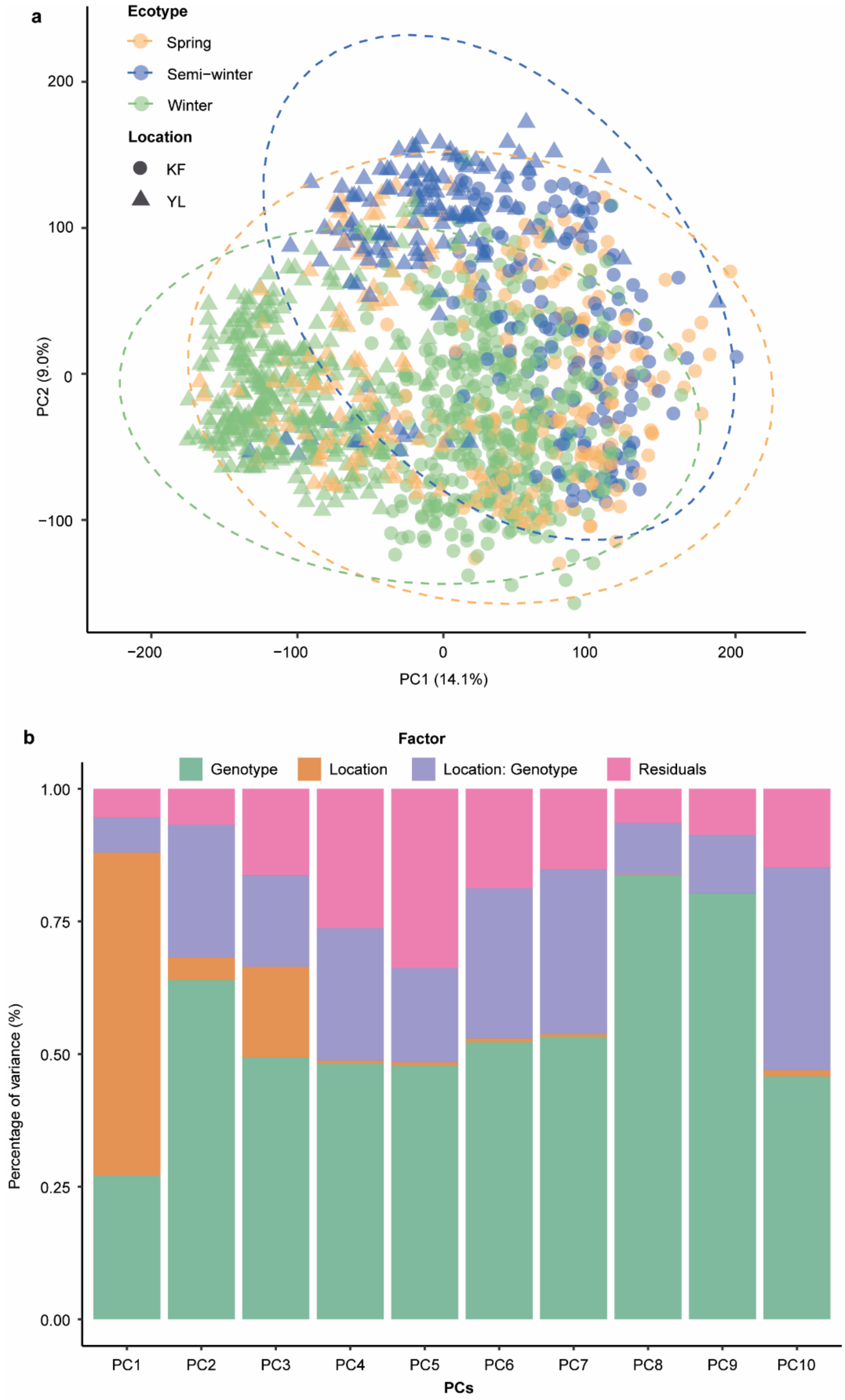


**Supplemental Figure 2. Overall transcriptome analysis for *B. napus* population (*n* = 175)**. (a) A principal component analysis illustrating the bacterial diversity across three different ecotypes at two locations. (b) Percentage of variance explained by genotype, location and location: genotype interaction.

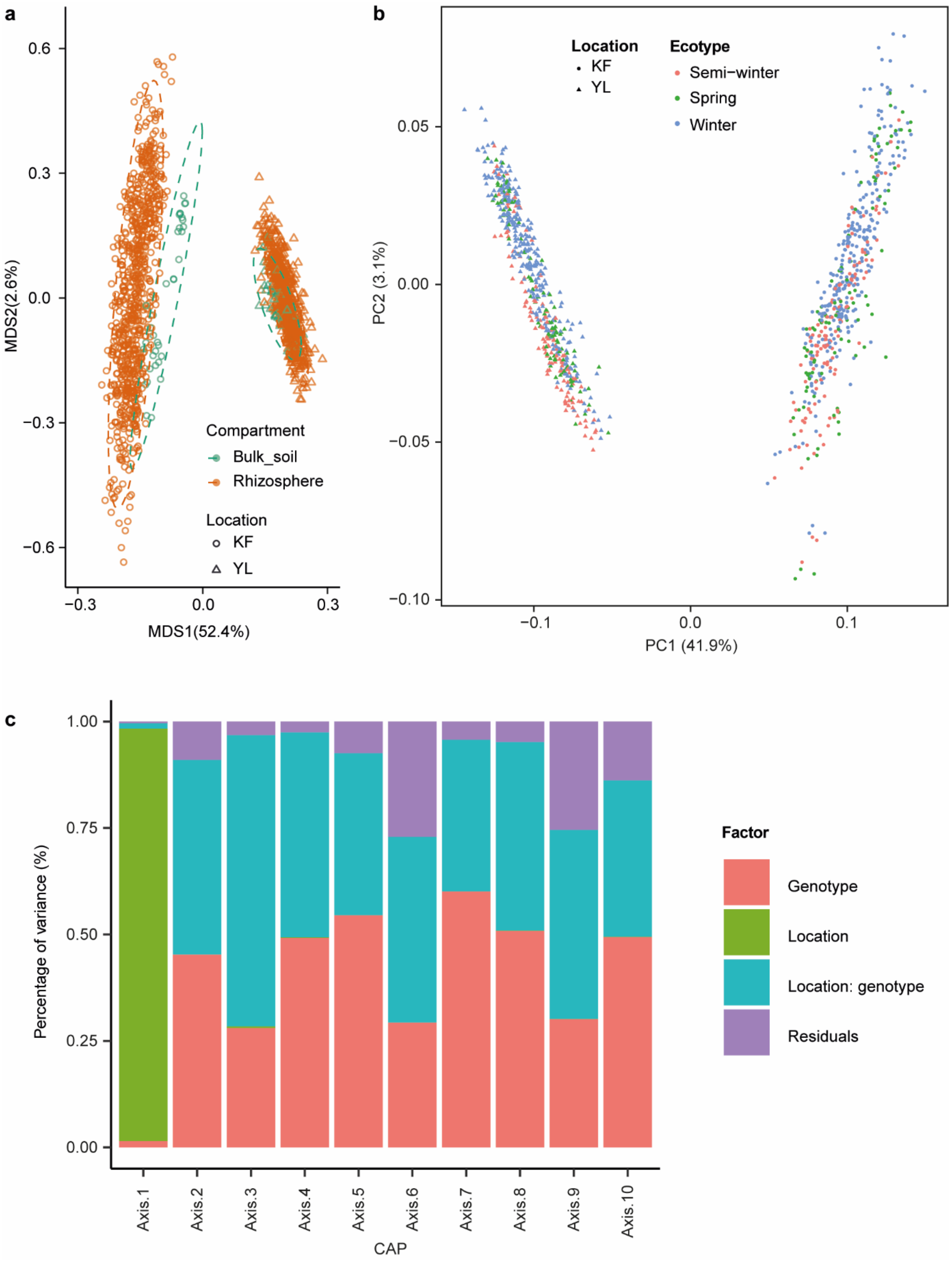


**Supplemental Figure 3. Large-scale bacterial microbiome profiling in the B. napus rhizosphere.** (a) A multidimensional scaling (MDS) plot illustrating the location and compartment effect on bacterial diversity. KF, Kaifeng; YL, Yangling. (b) A principal component analysis demonstrating the bacterial diversity across three different ecotypes at two locations. (c) Percentage of variance explained by genotype, location and location: genotype interaction.

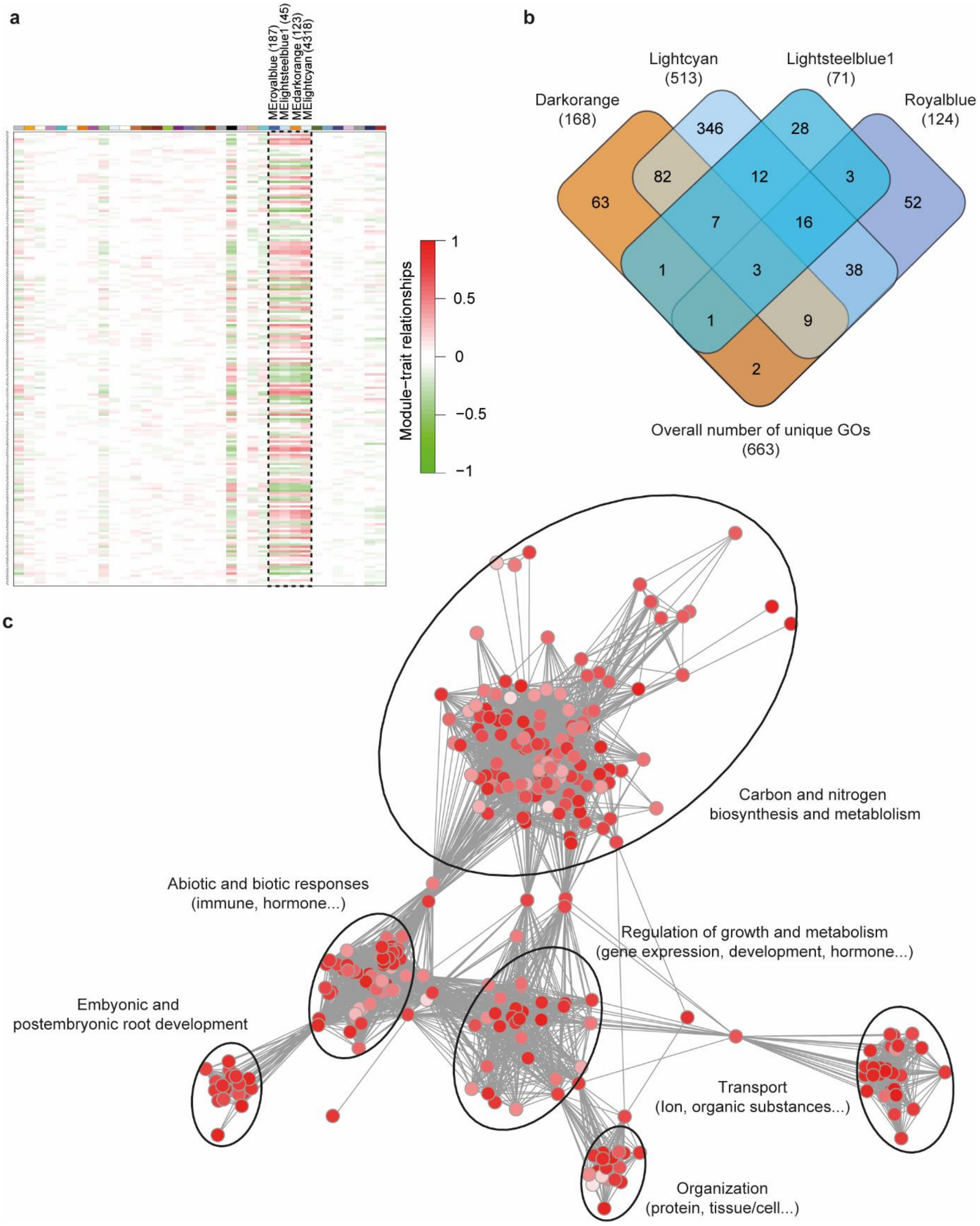


**Supplemental Figure 4. Identification of gene co-expressed modules and functional characterization.** (a) Weighted gene co-expression network analysis (WGCNA) identified 35 gene co-expressed modules. The number of genes enriched in each module was indicated in the brackets accordingly. The heatmap indicates the Pearson correlation of co-expressed modules with 203 heritable bacterial ASVs. (b) Overlapped unique Gene Ontology (GO) terms of four highly gene-microbial correlated modules. The number of GO terms enriched in each module was indicated in the brackets accordingly. (c) Network visualization of overlapped GO terms identified the major functional pathways enriched among the highly correlated WGCNA modules. The detailed pathways were annotated within the network associations done by REVIGO (http://revigo.irb.hr/).

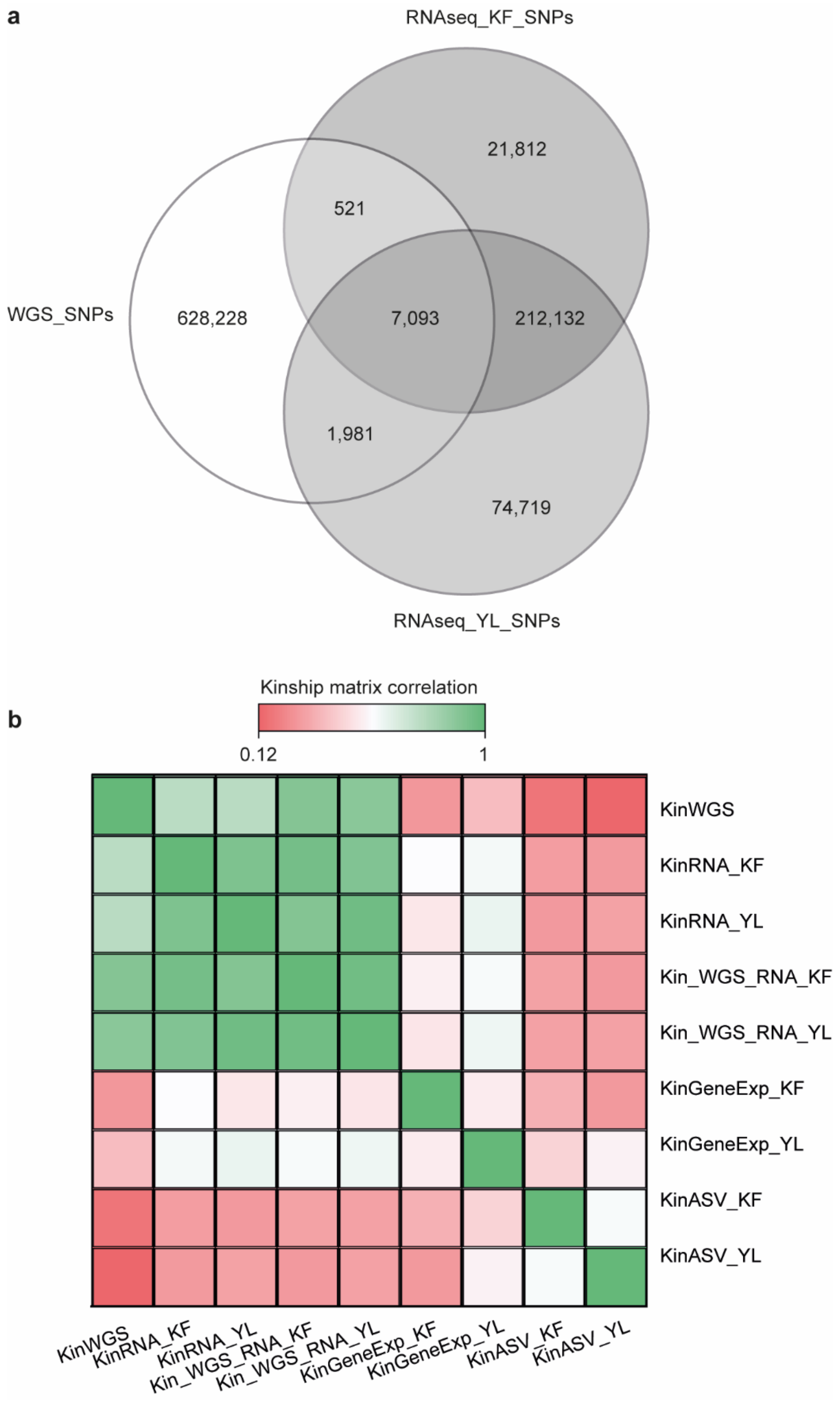


**Supplemental Figure 5. Preparation of multi-omics datasets used for genomic prediction.** (a) The overlapped SNPs among whole genome sequencing (WGS) SNPs and RNAseq derived SNPs from KF (Kaifeng) and YL (Yangling) locations respectively. (b) The correlation between different kinship matrix.

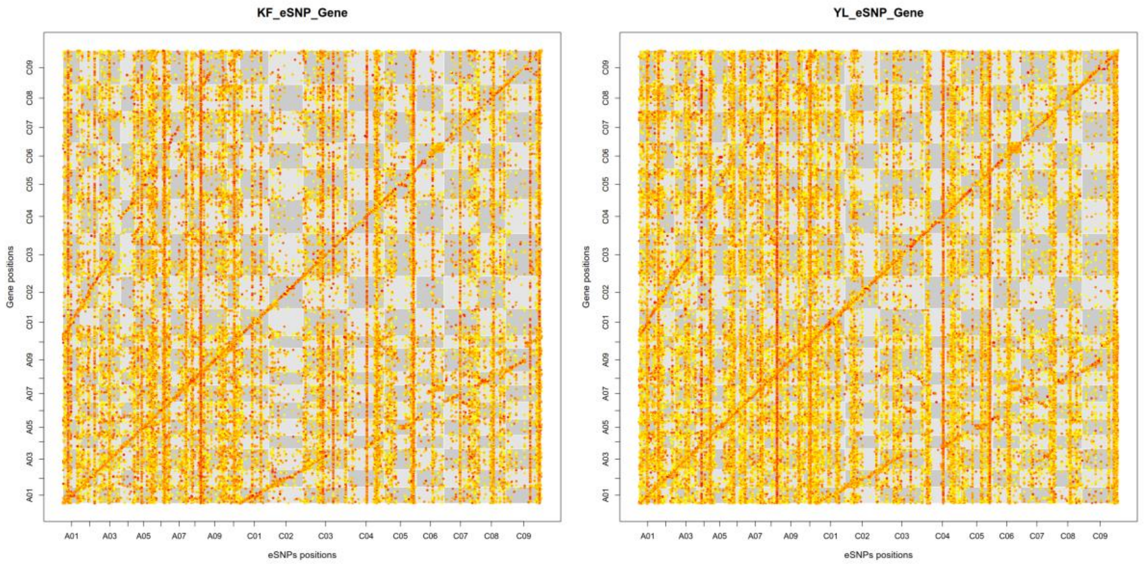


**Supplemental Figure 6. Dot plot showing eQTLs and their regulated genes in 19 chromosomes.** x-axis shows the single nucleotide polymorphism (SNP) position (bp) in each chromosome and the y-axis shows the gene position (bp) in each chromosome, with a chromosome order of A01 to C09 from left to right (x-axis) or from lower to upper (y-axis). The color of each dot represents the significance (P-value) of each eQTL-gene association, with low significance in yellow and high significance in red. Each chromosome is scaled by the physical chromosome length. Dots in the diagonal line show the intra-chromosomal associations. The lines show the enrichment of inter-subgenomic associations in homoeologous chromosomes and inter-chromosomal associations between the hotspot and other chromosomes.

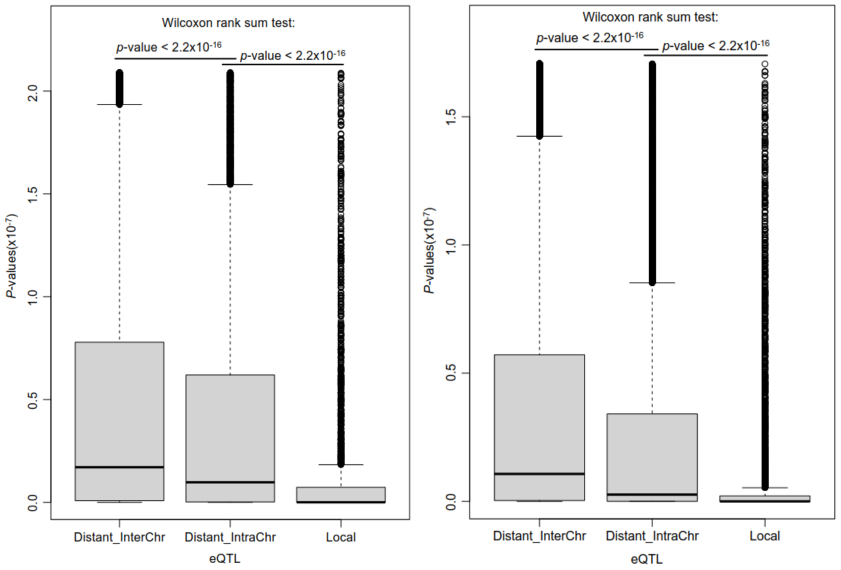


**Supplemental Figure 7. The comparison of significance (*P* values) between inter-chromosomal eQTLs (Distant_InterChr) and intra-chromosomal eQTLs (Distant_IntraChr and Local) in KF (left) and YL (right).**

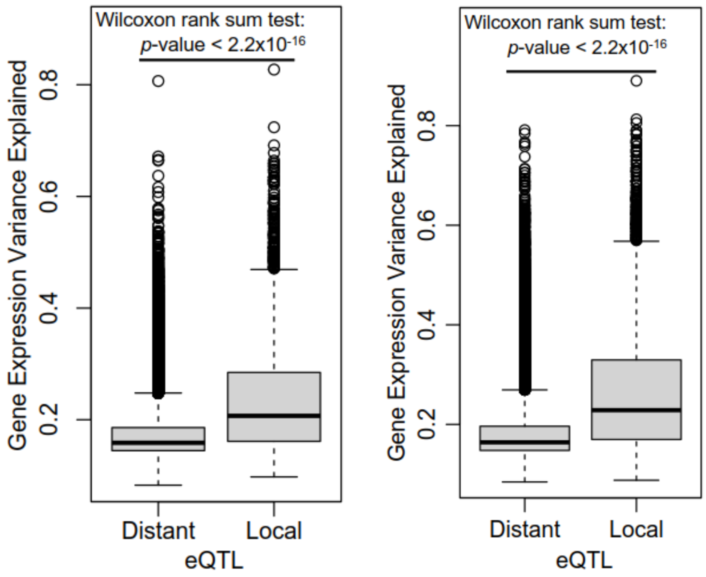


**Supplemental Figure 8. Difference of gene explanation variance explained (r2) of SNPs between local eQTL and distant eQTL in KF (left) and YL (right).**

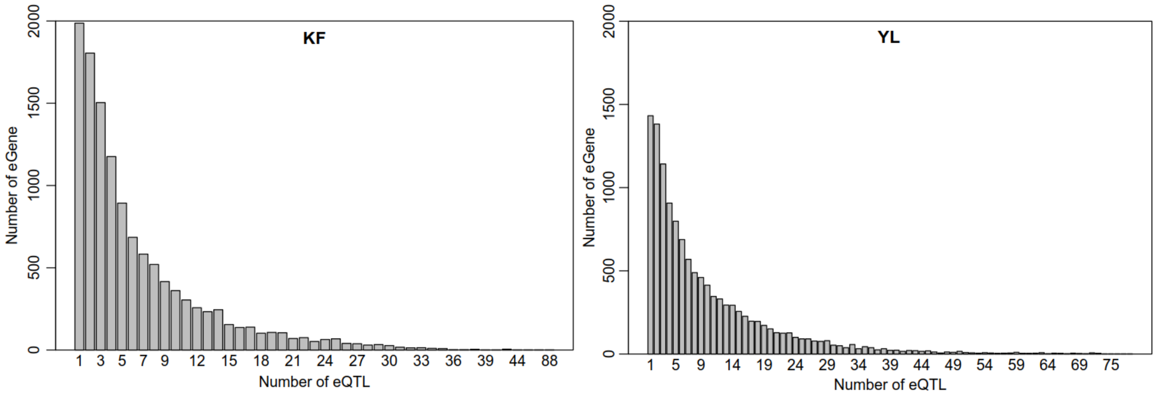


**Supplemental Figure 9. Distribution of the number of eQTLs for genes which were regulated by eQTL in KF (left) and YL (right).**

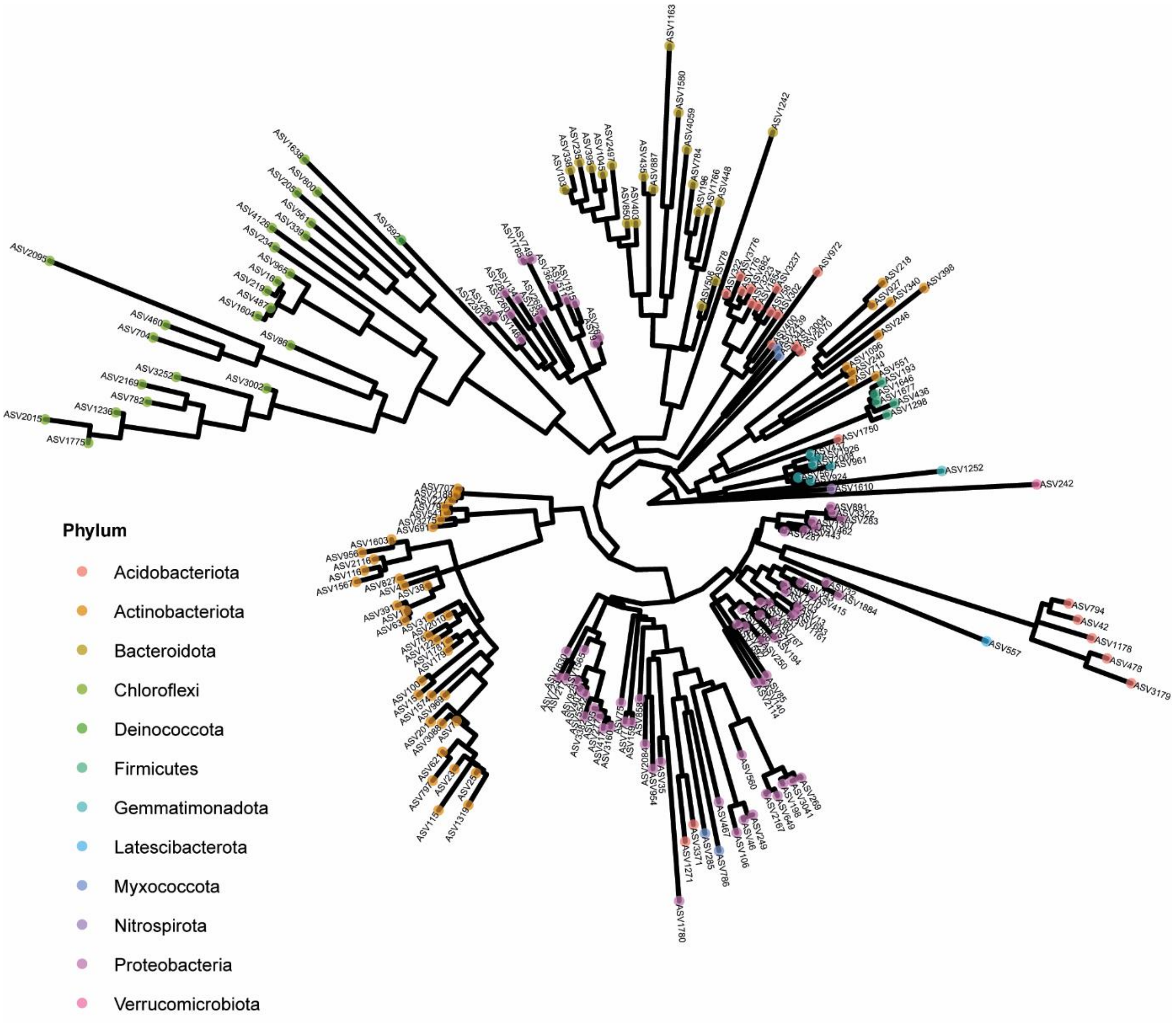


**Supplemental Figure 10. Phylogenetic tree of 203 highly heritable ASVs.** The complete list of 203 ASVs are provided in the Supplemental Dataset 3. Different taxonomic information were color coded at the family level.

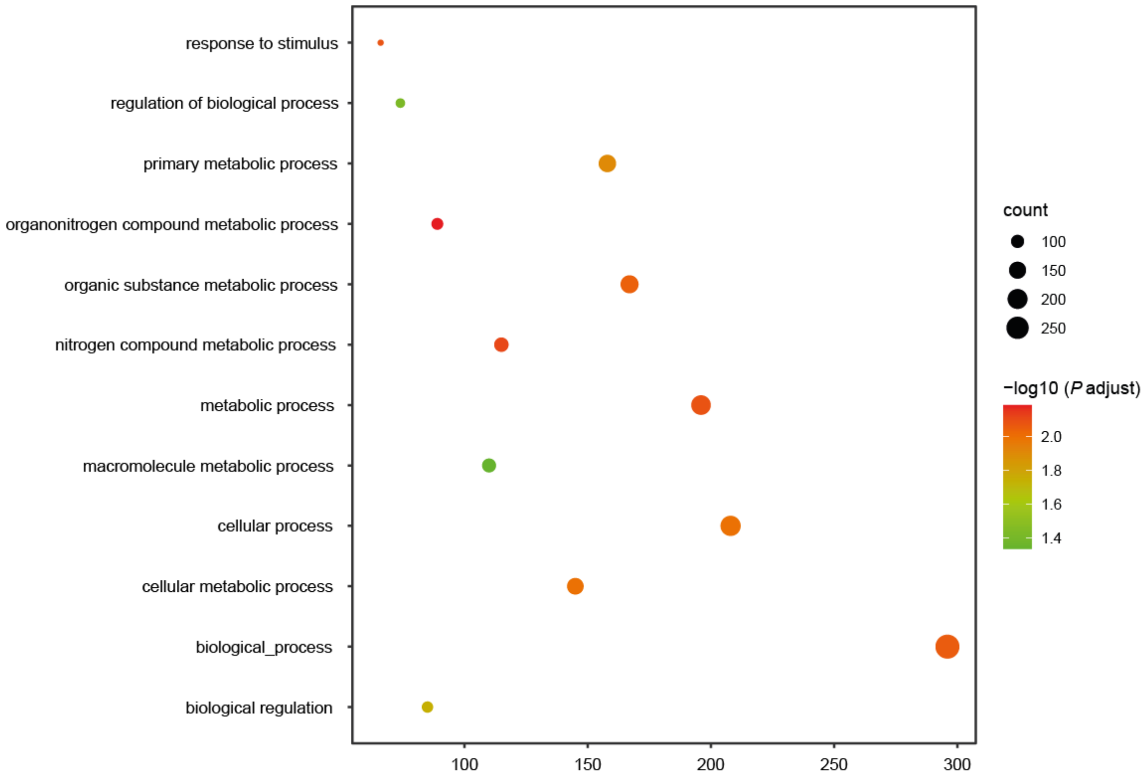


**Supplemental Figure 11. Gene ontology (GO) analysis of ASV729 associated genes.** The size of dots indicates the number of detected ASVs enriched into each biological process. The color code indicates the significance of enrichment.

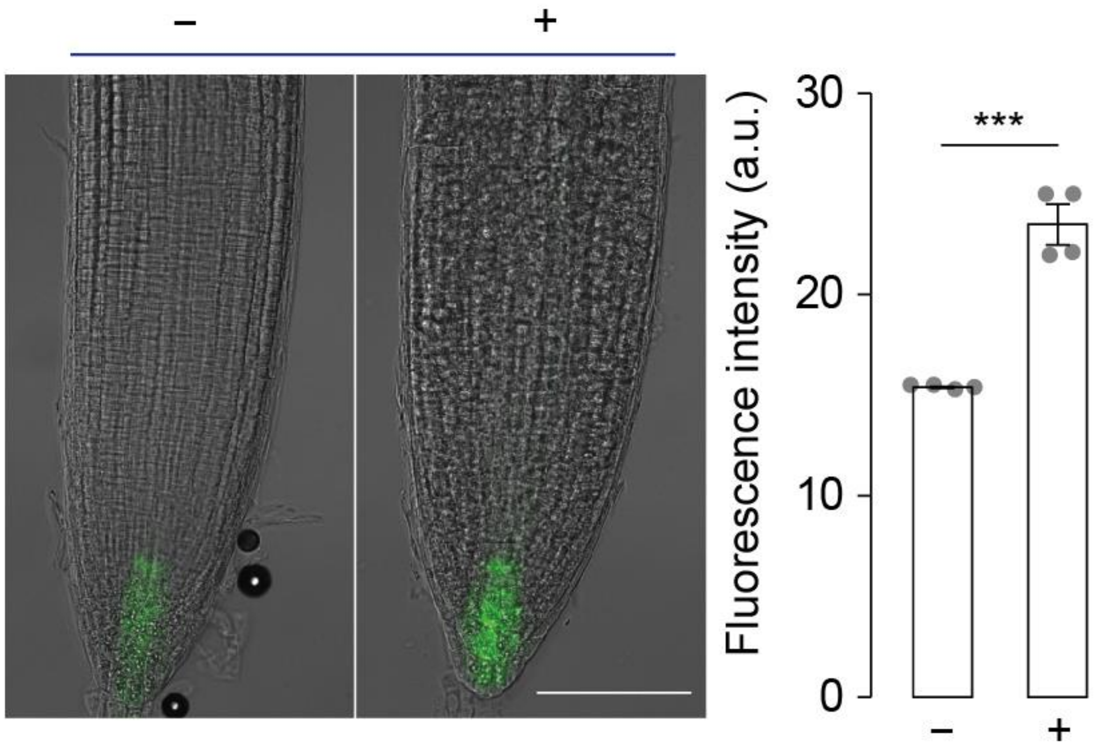


**Supplemental Figure 12. DR5::GUS tagged root tip imaging with or without inoculation of *Sphingopyxis* isolate (29-6000-31).** The significance was controlled by paired Student's *t* test. ***, *P* <0.001.

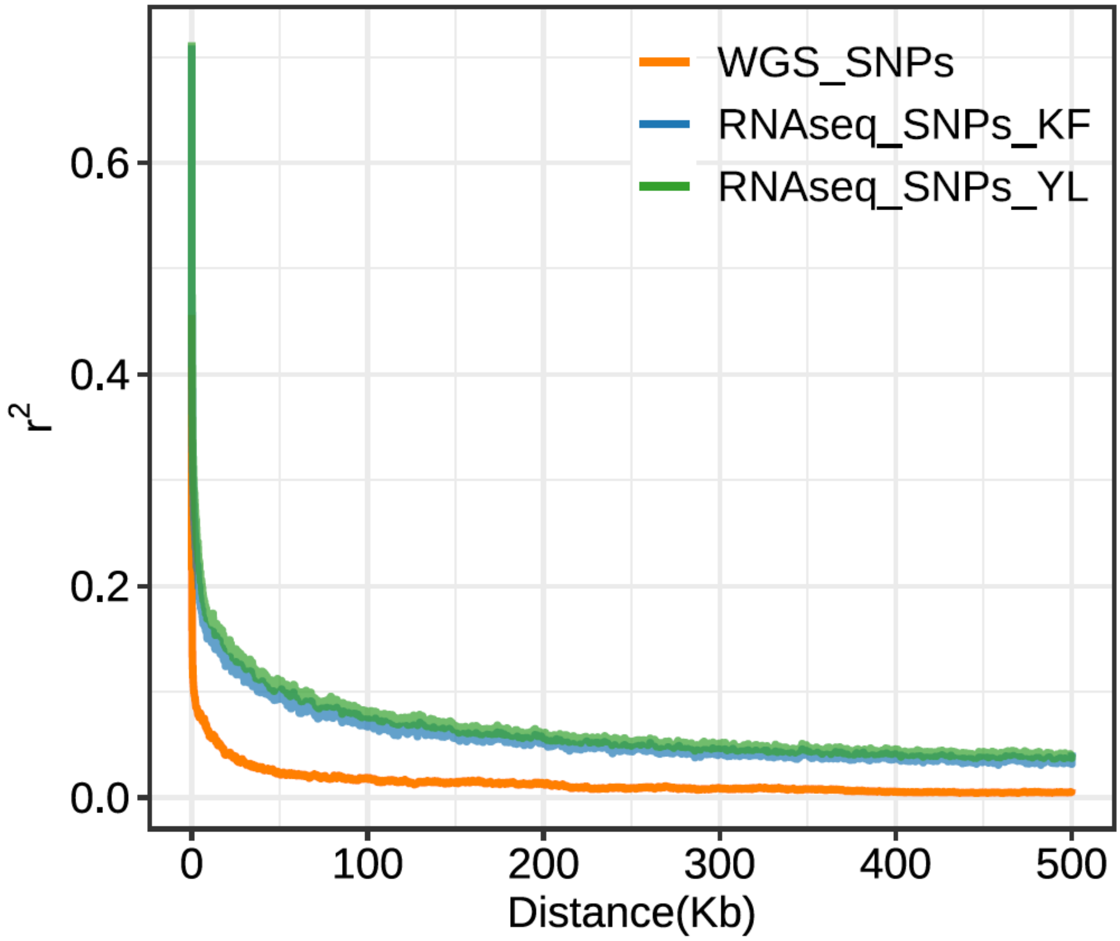


**Supplemental Figure 13. Linkage disequilibrium (LD) decay plot referring to the nonrandom associations of alleles at different loci for whole genome sequencing (WGS) and RNAseq derived SNPs data.** KF, Kaifeng; YL, Yangling.

**Supplemental Table 1 Physical and chemical properties of the basic soil at two locations.** The values presented are mean ± standard error at Kaifeng (KF) and Yangling (YL) fields in 2019. Means followed by the same letter are not significantly different at $p$ <0.05 according to LSD (ANOVA, turkey HSD, $n$ = 5).

| **Soil properties** | **Kaifeng (KF)** | **Yangling (YL)** |
|---|---|---|
| Total N (%) | 0.094 ± 0.0027a | 0.076 ± 0.00088b |
| Conc. P (mg/kg) | 69.23 ± 1.61a | 67.55 ± 1.06a |
| pH | 6.74 ± 0.08b | 7.23 ± 0.07a |
| Total organic carbon (%) | 0.60 ± 0.062a | 0.56 ± 0.043a |
